# Human assembloids reveal the consequences of *CACNA1G* gene variants in the thalamocortical pathway

**DOI:** 10.1101/2023.03.15.530726

**Authors:** Ji-il Kim, Yuki Miura, Min-Yin Li, Omer Revah, Sridhar Selvaraj, Fikri Birey, Xiangling Meng, Mayuri Vijay Thete, Sergey D. Pavlov, Jimena Andersen, Anca M. Pașca, Matthew H. Porteus, John R. Huguenard, Sergiu P. Pașca

## Abstract

Abnormalities in crosstalk between the thalamus and the cerebral cortex are thought to lead to severe neuropsychiatric disorders, such as epilepsy and psychotic disorders. Pathogenic variants in the CACNA1G gene, which encodes the α1G subunit of the thalamus-enriched T-type voltage-gated calcium channel CaV3.1, are associated with absence seizures, intellectual disability, and schizophrenia, but the cellular and circuit level consequences of these genetic variants in humans remain unknown. Here, we developed an in vitro human assembloid model of the thalamocortical pathway to systematically dissect the contribution of genetic variants in T-type calcium channels. We discovered that a CACNA1G variant (M1531V) associated with seizures led to changes in T-type currents in human thalamic neurons, as well as correlated hyperactivity of thalamic and cortical neurons in thalamo-cortical assembloids. In contrast, CACNA1G loss, which has been associated with risk of schizophrenia, resulted in abnormal thalamocortical connectivity that was related to both increased spontaneous thalamic activity and aberrant thalamic axonal projections. Taken together, these results illustrate the utility of organoid and assembloid systems for interrogating human genetic disease risk variants at both cellular and circuit level.

## Introduction

Communication between the thalamus and the cerebral cortex via reciprocal thalamocortical projections is essential for processing sensory information and for modulating cognitive functions in the mammalian nervous system (1-3). Anatomical or functional dysconnectivity in the thalamocortical circuit is often associated with psychiatric disorders, including schizophrenia and autism spectrum disorder (4, 5). Moreover, hypersynchronous thalamocortical activity is a core feature of absence seizures in epilepsy (6, 7). The thala-mus-enriched voltage-dependent calcium channel Ca_V_3.1, encoded by the *CACNA1G* gene, is a key mediator of activity in this circuit (8-11). Importantly, pathogenic variants in *CACNA1G* are strongly associated with disease; putative loss-of-function variants have been linked to schizophrenia and intellectual disability (12, 13), while putative gain-of-function variants have been linked to epilepsy (14-16). However, the cell-specific consequences of different human *CACNA1G* gene variants are unknown. Further, how these cellular defects influence activity in the thalamocortical pathway is a mystery. Addressing this has been particularly challenging in the context of evolutionary changes in thalamocortical projections in primates, in which, for instance, thalamocortical projections innervate the dorsal forebrain at earlier stages than in rodents (17).

We previously introduced an assembloid approach to model network-level interactions in the developing nervous system by functionally integrating regionalized neural organoids (18, 19), and applied these models to study disease mechanisms (20-23). Despite efforts to derive thalamus-like neural cells from mouse embryonic stem cells (24) and human induced pluripotent stem (hiPS) cells (25) and to assemble them with cortical neurons, it remains unclear if these in vitro neurons recapitulate distinctive electrophysiological properties of thalamic neurons, such as T-type calcium channel-dependent rebound firing, and whether they can functionally connect in assembloids. For instance, while previous work showed that, in principle, thalamus organoids can send axonal projections into *ex vivo* mouse cortical tissue (24) or into human cortical organoids (25), functional connectivity through concomitant electrophysiological recordings and following functional manipulation has not been demonstrated. Moreover, the range of dynamic responses and variability required for disease modeling has not been established for thalamo-cortical assembloids, and how closely these thalamic cultures resemble the diencephalon has been precluded by lack of transcriptomic data in primary human tissue.

Here, we developed a platform that combines assembloids, live cell imaging, optogenetics, and extracellular recordings to investigate early development of the human thalamocortical pathway. Using hiPS cells, we generated human 3D regionalized neural organoids resembling the diencephalon, the developmental precursor to thalamic glutamatergic neurons. The neurons in human diencephalic organoids (hDiO) transcriptionally and electrophysiologically recapitulated features of thalamic glutamatergic neurons. When assembled with human cortical organoids (hCO), these neurons sent thalamocortical projections to the cortical neurons. Using optogenetics and simultaneous extracellular recordings or calcium imaging, we demonstrated the fomation of the functional thalamocortical pathway in these assembloids. To study *CACNA1G*-related channelopathies, we generated hiPS cell lines carrying heterozygous and homozygous gain- and loss-of-function variants in *CACNA1G*. Probing thalamo-cortical assembloids revealed potential defects in T-type current decay and correlated hyperactivity in the gain-of-function *CACNA1G* variant; in contrast, *CACNA1G* loss-of-function resulted in hyperactivity following abnormal thalamocortical connectivity.

## Results

### Generation of 3D human diencephalic organoids (hDiO) from hiPS cells

To generate neural organoids that include thalamic glutamatergic neurons, we implemented guided differentiation approach to generate regionalized neural organoids from hiPS cells, as we have done previously for other nervous system domains (20, 21, 26, 27). We applied small molecules that modulate the SHH (SAG), WNT (CHIR), and BMP (BMP7) signaling to differentiate cells to diencephalic-like fates (Fig. 1A, Supplementary Fig. 1A). These morphogens are known to control the development of the early diencephalon and thalamus (28-32). Of note, previous approaches to generate thalamic neurons from mouse stem cells or hiPS cells used BMP7, insulin, and PD0325901 (24, 25). The neural organoids treated with a combination of 1 µM CHIR, 100 nM SAG and 30 ng/mL of BMP7 showed a higher expression of the diencephalic markers *TCF7L2* and *OTX2* and a lower expression of the rostral forebrain marker *FOXG1* when compared to human cortical organoids (hCO) (Supplementary Fig. 1B). We refer to these regionalized organoids with the broader term human diencephalic organoids (hDiO) (33). To examine cellular diversity more comprehensively in these organoids, we performed droplet-based single-cell RNA sequencing (scRNAseq) at day 100 of differentiation (Fig. 1A, Supplementary Fig. 1C). Uniform Manifold Approximation and Projection (UMAP) showed the presence of a large cluster of glutamatergic neurons that express thalamic glutamatergic neuronal markers *TCF7L2* and *SLC17A6*. This analysis also revealed clusters of GABAergic neurons (*SLC32A1*), neuronal progenitors (*GADD45G*), choroid plexus-like cells (*TTR*), astrocytes (*AQP4*), OLIG2-expressing cells, proliferating cells (*MKI67*), and a negligible number of cells expressing the rostral forebrain marker *FOXG1* or the dopaminergic neuron marker *TH* (Fig. 1B, Supplementary Fig. 1D). We observed choroid plexus-related cells in our hDiOs and this may be related to BMP7 signaling as BMP4 is known to induce choroid plexus-like cells (34, 35). Unbiased mapping of scRNAseq data onto midgestational human samples using VoxHunt demonstrated hDiO approximates post-conceptional week (PCW) 15–24 (36, 37) with high similarity to the human mediodorsal nucleus of the thalamus (MD) (Fig. 1C, Supplementary Fig. 1E). In addition, VoxHunt analysis using mouse in situ gene expression data indicated high scaled correlation of hDiO with the diencephalon of embryonic day 15.5 mouse (Fig. 1D). We also confirmed the presence of cells in the hDiO expressing early developing thalamic regional markers (38) for caudal thalamus (cTh) (*GBX2, CD47*), rostral thalamus (rTH) (*SOX14, GATA3*), and epithalamus (Eth) (*TAC1, POU4F*), with neglectable expression of zona limitans intrathalamica (ZLI) (*PITX2*, FOXA2) and pretectum (rPT) (*1C1QL2, BHLHE23*) markers (Supplementary Fig. 1F). To further validate the transcriptional similarity between cells in vitro and their in vivo counterpart, we compared scRNAseq data from hDiO and the developing human thalamus (Fig. 1E–G, Supplementary Fig. 1G). We obtained a PCW20 diencephalic region, performed scRNAseq analysis, and combined this with the publicly available PCW16–20 human thalamus scRNAseq data (39). UMAP visualization demonstrated that thalamic glutamatergic neurons (*SLC17A6*), GABAergic neurons (*GAD1*), astrocytes (*GFAP*), glial progenitors (*SOX9*), oligodendrocytes (*SOX10*) and cycling cells (*MKI67*) are present in both hDiO and primary tissues. In contrast, endothelial cells (*FLT1*), microglia (*CSF1R*), mesenchymal cells (*COL1A1*), pericytes (*HIGD1*), blood cells (*HBA1*), and melanocytes (*DCT*) were only detected in primary tissue, while choroid plexus-like cells (*TTR*) were only found in hDiO (Fig. 1E, Supplementary Fig. 1G). We found that the proportion of *TCF7L2*^+^ cells in day 100 hDiO (∼60%) was within the range observed in primary tissue at PCW16–20 (46–72%). Thalamic glutamatergic neurons (*TCF7L2*^+^/*SLC17A6*^+^/*NTNG1*^+^) overlapped well across datasets (Fig. 1F) and represented approximately up to 45% of cells in hDiO and primary tissues over time (Fig. 1G). To further examine expression of thalamic glutamatergic neuronal markers at protein levels in hDiO, we performed immunostaining for *TCF7L2*, which was highly expressed in human diencephalon tissue at PCW19 (Fig. 1H). We found an increased expression of TCF7L2 in hDiO compared to hCO (Fig. 1I). Additionally, approximately 80% of the neurons labeled with a neuron-specific promoter (AAV-hSYN1::EYFP) in hDiO were VGLUT2^+^ neurons indicating that a predominant subset of neurons in hDiOs have glutamatergic identity (Fig. 1J). Lastly, we characterized the electrophysiological properties of hDiO neurons. Patch-clamp recordings from AAV-hSYN1::EYFP labeled hDiO neurons showed spontaneous action potentials (Supplementary Fig. 2A), as well as characteristic post-hyperpolarization rebound spiking (40-42) (Fig. 1K, Supplementary Fig. 2B). The rebound spikes were blocked by the T-type calcium channel blocker TTA-P2 (Fig. 1K). The current-voltage (I-V) curve (measured by barium currents) showed a typical low-voltage activation profile, with activation voltage threshold around –50 mV and a peak current between –30 mV and –20 mV (Supplementary Fig. 2C). The fast T-type channel-dependent calcium influx was measured by barium current recordings, which could be blocked by TTA-P2 (43, 44) (Supplementary Fig. 2D, E). T-type currents also showed high reliability for repetitive trials (Supplementary Fig. 2D). In summary, neurons in hDiO transcriptionally and electrophysiologically resemble thalamic glutamatergic neurons.

**Figure 1.**
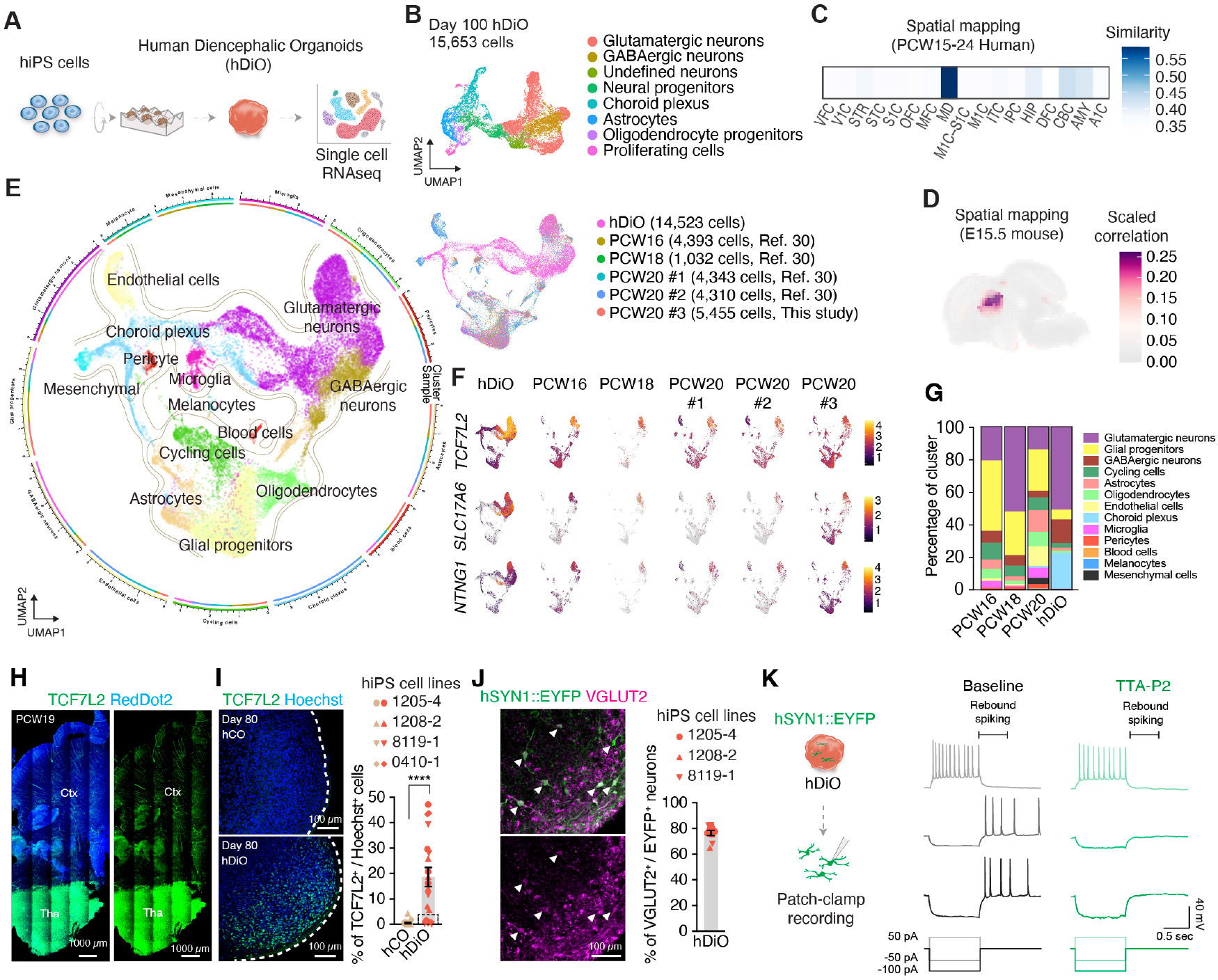
Generation of 3D human diencephalic organoids (hDiO). (**A**) Schematic describing the generation of 3D human diencephalic organoids (hDiO) from hiPS cells and scRNAseq analysis. (**B**) UMAP visualization of single cell RNA expression in hDiO at day 100 of *in vitro* differentiation (*n* = 15,653 cells from 3 hiPS cell lines). (**C**) VoxHunt spatial brain mapping of hDiO onto the BrainSpan dataset of the human developing brain (PCW15– 24). (**D**) VoxHunt spatial brain mapping of hDiO onto the Allen Brain Institute E15.5 mouse brain data. (**E**) UMAP of integrated hDiO and human primary samples from PCW16–20. (**F**) UMAP visualization of thalamic glutamatergic markers *TCF7L2, SLC17A6* and *NTNG1*, and (**G**) percentage distribution of cell clusters in each sample. (**H**) CUBIC-cleared human primary samples immunostained with TCF7L2 (green) and RedDot2 nuclear marker (magenta). Scale bar: 1000 μm. (**I**) Immunostaining for TCF7L2 (green), and Hoechst (blue) in hCO and hDiO at day 80, and quantification of TCF7L2^+^ cells (*n* = 20 for hCO and *n* = 19 for hDiO from 3 differentiation experiments of 4 hiPS cell lines; Mann-Whitney test, \*\*\*\**P* < 0.0001). Scale bar: 100 μm. We noted that several organoids had very low levels of TCF7L2 expression, and these came from one differentiation batch (dashed box), which we deemed unsuccessful and not used organoids in subsequent assays. (**J**) Immunostaining for GFP (green) and VGLUT2 (magenta) in hDiO at day 80, and quantification of VGLUT2^+^/EYFP^+^ cells (*n* = 18 from 2 differentiation experiments of 3 hiPS cell lines). Scale bar: 100 μm. (**K**) Schematic describing whole-cell patch-clamp recording of hDiO neurons (left) and representative traces of hyperpolarization-induced rebound spikes (500 ms window) in AAV-hSYN1:: EYFP^+^ neurons (right). Cells were held at –50 mV for rebound spike recordings and were held at –60 mV for depolarization-induced spikes. Examples shown for –100 pA and –50 pA current induced rebound spikes. Rebound spikes were blocked by bath application of TTA-P2 (3 μM). Data is shown as mean ± s.e.m. Dots indicate organoids in I, J and cells in L. Each shape represents a hiPS cell line: Circle, 1205-4; Triangle, 1208-2; Inverted triangle: 8119-1; Rhombus, 0410-1 in I, J. Abbreviations: VFC; ventrolateral frontal cortex, V1C; primary visual cortex, STR; striatum, STC; Posterior superior temporal cortex, S1C; primary somatosensory cortex, OFC; orbital frontal cortex, MFC; medial frontal cortex, MD; mediodorsal nucleus of thalamus, M1C; primary motor cortex, ITC; inferior temporal cortex, IPC; posterior inferior parietal cortex, HIP; hippocampus, DFC; dorsolateral prefrontal cortex, CBC; cerebellar cortex, AMY; amygdala, A1C; primary auditory temporal cortex.

### Projections and functional connectivity in thalamo-cortical assembloids

To model the formation of thalamocortical connectivity in vitro, we used an assembloid approach as we have previously described for other regionalized neural organoids (21, 45) (Fig. 2A). We virally labeled the hCO with hSYN1:: mCherry and the hDiO with hSYN1::EYFP and then assembled them by placing the two regionalized organoids in close contact in an Eppendorf tube. After 72 hours, the two organoids fused to generate a thalamo-cortical assembloid (Fig. 2B). To quantify the structural integration between the two organoids, we measured the fluorescence area on each side of the assembloid. We found a progressive increase in reciprocal, bilateral projections between hCO and hDiO over daf 30 (days after fusion) (Fig. 2C, D). This is consistent with early embryonic stage formation of thalamocortical and cortico-thalamic projections in primate embryos (17) and in contrast with the formation of unilateral projections in cortico-striatal assembloids (21). To verify connections between hDiO projection neurons and hCO neurons, we applied anterograde tracing (46) by expressing AAV1-hSYN1::Cre in hDiO and AAV-EF1α::DIO-tdTomato in hCO (Fig. 2E). At daf 36, we found EYFP^+^/tdTomato^+^ cells on the hCO side of the thalamo-cortical assembloids, suggestive of thalamo-cortical connectivity (Fig. 2F). We next used retrograde tracing by infecting hCO with rabies-ΔG-Cre-EGFP and AAV1-EF1α::rabies glycoprotein (G) and hDiO with AAV-EF1α::DIO-tdTomato (47, 48) before generating thalamo-cortical assembloids (Fig. 2G). At daf 24, we observed EGFP^+^/tdTomato^+^ double-positive cells on the hDiO side of thalamo-cortical assembloids (Fig. 2H). These experiments suggest anterograde and retrograde transmission in assembloids through thalamocortical connections.

**Figure 2.**
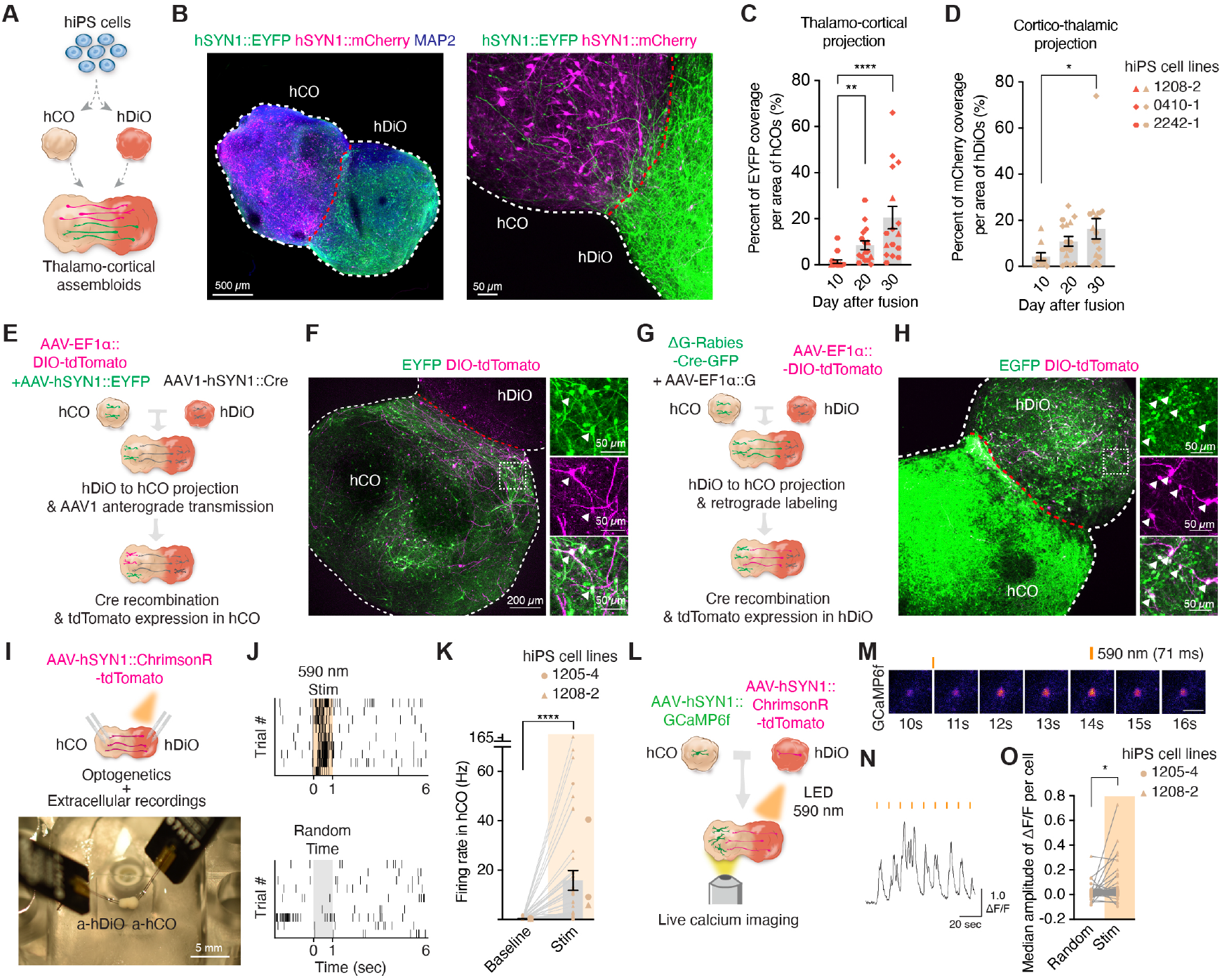
Generation of human assembloids to study thalamocortical connectivity. (**A**) Schematic illustrating the thalamocortical pathway in assembloid model. (**B**) Representative images of a 3D thalamo-cortical assembloid showing an entire assembloid (left) and thalamocortical projections at high magnification (right). (**C, D**) Quantification of EYFP and mCherry projections in assembloids at daf 10, 20 and 30. (*n* = 10–16 assembloids from 4 differentiation experiments of 3 hiPS cell lines; Kruskal-Wallis test, \**P* = 0.0273, **P = 0.0028, \*\*\*\**P* < 0.0001). (**E**) Schematic of anterograde viral tracing in thalamo-cortical assembloids using AAV1. (**F**) Representative image showing the presence of EYFP^+^/tdTomato^+^ cells in the a-hCO side of thalamo-cortical assembloids at daf 36. (**G**) Schematic illustrating retrograde viral tracing in thalamo-cortical assembloids using rabies virus. (**H**) Representative image indicating EGFP^+^/tdTomato^+^ cells in a-hCO at daf 24. (**I**) Schematic illustrating extracellular recording coupled with optogenetic stimulation of thalamo-cortical assembloids (top). Image showing insertion of each electrode into a-hCO and a-hDiO in thalamo-cortical assembloids (bottom). (**J**) Raster plot showing optically-evoked responses in thalamo-cortical assembloids (top) and randomly-selected time points (bottom).(**K**) Quantification of firing rates in a-hCO in assembloids at daf 56–91 before and after 590 nm light application (*n* = 49 units from 5 assembloids, 3 differentiation, 2 hiPS cell lines; Wilcoxon test, \*\*\*\**P* < 0.0001). (**L**) Schematic illustrating calcium imaging coupled with optogenetics in thalamocortical assembloids. (**M**) Representative images and (**N**) calcium imaging trace of cells in a-hCO following light stimulation of a-hDiO. (**O**) Median amplitude of ΔF/F signals of optically-evoked responses on the a-hCO side of thalamo-cortical assembloids at daf 45–54 (*n* = 176 cells from 2 differentiation experiments of 2 hiPS cell lines; Wilcoxon test, \**P* = 0.039). Dots indicate assembloids in C, D. Small dots indicate units or neurons in K, O and large dots indicate assembloids in K. Each shape represents a hiPS cell line: Circle, 1205-4; Triangle, 1208-2; Hexagon, 2242-1 in C, D, K, O. Abbreviations: Daf; days after fusion of assembloids.

To examine whether thalamo-cortical assembloids form functional connections, we combined optogenetic stimulation with simultaneous extracellular recording in thalamo-cortical assembloids using a custom extracellular recording system (Fig. 2I, Supplementary Fig. 3A). Acute recordings from individual hDiO using Cambridge NeuroTech^TM^ ultrathin and high-density electrode arrays (Supplementary Fig. 3B) revealed spontaneous neuronal activity (Supplementary Fig. 3C). Optogenetic stimulation of hDiO expressing hSYN1::ChrimsonR–tdTomato (49) elicited increased firing of isolated single units (Supplementary Fig. 3D,E), which was blocked by sodium channel blocker tetrodotoxin (TTX) (Supplementary Fig. 3E,F). To probe functional connectivity in thalamo-cortical assembloids, we optogenetically stimulated hDiO of the assembloid (a-hDiO) while recording extracellular signals from the hCO side of the assembloid (a-hCO) (Fig. 2I–K). We found increased firing rates of single units in a-hCO following light illumination (Fig. 2J, K). In contrast, light illumination did not influence firing rates in a-hCO when the opsin was not expressed in a-hDiO (Supplementary Fig. 3G–I), demonstrating that optically-evoked responses in the thalamo-cortical assembloids are not due to light-related artifacts. We also delivered AAV-hSYN1::GCaMP6f to a-hCO and AAV-hSYN1::ChrimsonR-tdTomato to a-hDiO, and then applied 590 nm light and measured calcium activity in a-hCO neurons after thalamic stimulation (Fig. 2L). Consistent with extracellular recording experiments, we detected significantly increased light-dependent calcium responses in a-hCO neurons in thalamo-cortical assembloids (Fig. 2M–O). Altogether, these results support the presence of anatomical and functional connectivity in hiPS cell-derived thalamo-cortical assembloids.

### Cellular defects in thalamic neurons carrying disease variants in the ***CACNA1G*** gene

The low-threshold voltage-dependent calcium channel Ca_V_3.1, encoded by the *CACNA1G* gene, is highly expressed in thalamic neurons in primary human fetal tissue, as well as in hDiO (Supplementary Fig. 4A, B). Both presumed gain- and loss-of-function variants in *CACNA1G* lead to severe channelopathies (12-16), but the consequences of these variants on human thalamic neurons and on their cortical neuronal targets remain unknown.

We generated hiPS cell lines with gain- and loss-of-function mutations in the *CACNA1G* gene using the 1208-2 hiPS cell line (Fig. 3A). To derive a homozygous gain-of-function line, we used AAV6/Cas9 RNP genome-editing methods in hiPS cells (50) resulting in a substitution of methionine 1531 to valine in domain III of *CACNA1G* (*CACNA1G* M1531V; Fig. 3B, C, Supplementary Fig. 4C-E). De novo heterozygous *CACNA1G* M1531V variant was reported in patients with early onset seizures and epileptic encephalopathy (14, 15).

**Figure 3.**
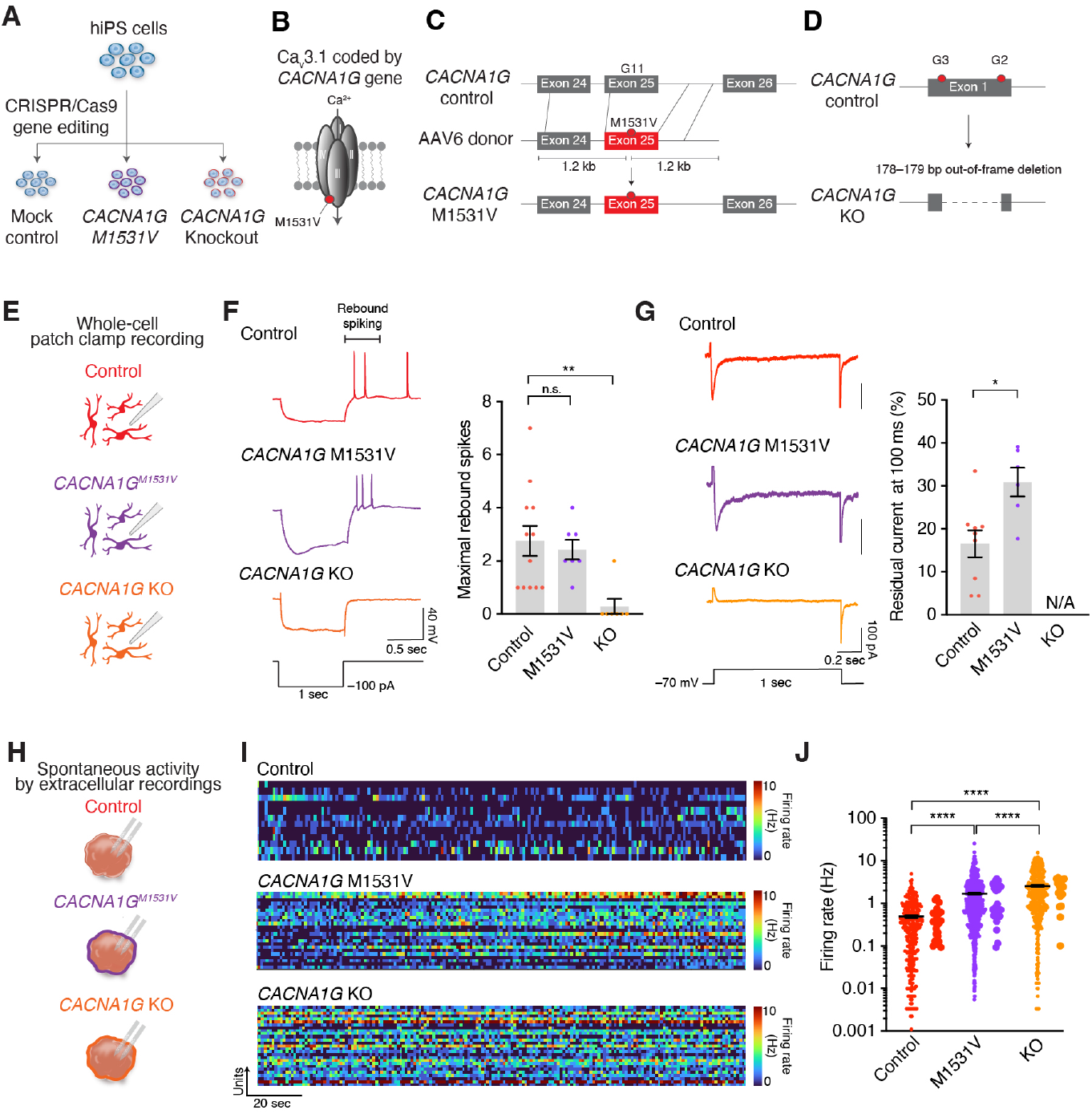
Neuronal activity in hDiO carrying pathogenic variants in the *CACNA1G* gene. (**A**) Schematic illustrating the generation of *CACNA1G* M1531V and KO hiPS cells using CRISPR/Cas9 genome editing. (**B**) Schematic illustrating the position of the M1531V variant in Ca_V_3.1. (**C**) Schematic illustrating the strategy to introduce the *CACNA1G* M1531V point mutation. (**D**) Schematic illustrating the strategy to introduce homozygous *CACNA1G* KO via exon 1 deletion. (**E**) Schematic illustrating whole-cell patch clamp recording of hDiO neurons from control and *CACNA1G* variants hiPS cell lines. (**F**) Representative traces showing hyperpolarization-induced rebound spikes in control, *CACNA1G* M1531V and *CACNA1G* KO hDiO neurons (left) and quantification of the number of maximal rebound spikes within the time window of 500 ms post-hyperpolarization (right) (n = 12 neurons for control, n = 7 neurons for *CACNA1G* M1531V and n = 8 neurons for *CACNA1G* KO; Kruskal-Wallis test, \*\**P* = 0.0014). (**G**) Representative fast T-type current traces induced by depolarization (from –70 mV) from control, *CACNA1G* M1531V and *CACNA1G* KO hDiO neurons (left) and quantification of the residual current at 100 ms after peak (right) (n = 9 neurons for control, n = 6 neurons for *CACNA1G* M1531V, not-applicable (N/A) for *CACNA1G* KO; Mann-Whitney test, \**P* = 0.0176). Cells were held at –70 mV, and voltage steps from –90 mV to +20 mV were applied. Fast T-type currents were measured at –30 mV. (**H**) Schematic illustrating the experimental design for probing spontaneous activity via extracellular recording from control, *CACNA1G* M1531V, and *CACNA1G* KO hDiOs. (**I**) Representative heatmaps of spontaneous extracellular recording from control (top), *CACNA1G* M1531V (middle) and *CACNA1G* KO (bottom) hDiOs. (**J**) Firing rates of sorted single units from control, *CACNA1G* M1531V, and *CACNA1G* KO hDiOs at day 120–182 (n = 273 units from 23 hDiOs, 5 differentiations for control, n = 641 units from 26 hDiOs, 6 differentiations for *CACNA1G* M1531V and n = 350 units from 17 hDiOs, 5 differentiations for *CACNA1G* KO; Kruskal-Wallis test, \*\*\*\**P* < 0.0001). Data is shown as mean ± s.e.m. Dots indicate cells in F, G. Small dots indicate units and large dots indicate assembloids in J.

To generate homozygous *CACNA1G* knockout (KO, *CACNA1G*^−/−^) hiPS cell lines, we used dual guide RNAs (gRNAs) targeting the exon 1 region of *CACNA1G* (Fig. 3D). We found that gene editing with a combination of 2 gRNAs efficiently caused 178 or 179 bp deletions in exon 1 (Supplementary Fig. 4F–I). We used 2 KO hiPS cell lines for subsequent assays. The engineered hiPS cell lines maintained their pluripotency (Supplementary Fig. 4J) and could be differentiated into regionalized organoids (Supplementary Fig. 4K). To verify whether *CACNA1G* variants influence patterning or cell composition of hDiOs, we performed scRNA-seq of hDiOs derived from mutant lines as well as the control 1208-2 hiPS cell line and found no major differences (Supplementary Fig. 5A, B). These samples also mapped to the E15.5 thalamus by VoxHunt analysis (Supplementary Fig. 5C). In addition, hDiOs had significantly more TCF7L2^+^ cells than hCOs across all hiPS cell lines (Supplementary Fig. 5D, E). This suggests that loss of *CACNA1G* and the M1531V gene variant do not result in major hDiO cellular composition changes.

To investigate how the *CACNA1G* pathogenic variants influence Ca_V_3.1 function in human thalamic neurons, we performed patch-clamp recordings of hDiO neurons (Fig. 3E). We measured rebound spikes after hyperpolarization, which require Ca_V_3.1 channels. We observed a significant reduction in rebound spikes in *CACNA1G* knockout neurons, while rebound spikes in *CACNA1G* M1531V neurons were comparable to controls (Fig. 3F). Barium currents, which are a proxy of calcium channel function, were also significantly reduced in *CACNA1G* KO hDiO neurons (Supplementary Fig. 6J). Moreover, we looked for evidence of prolonged inactivation of T-type currents in neurons carrying the *CACNA1G* M1531V mutation by measuring the amount of residual T-type current at 100 ms after the peak when Ca_V_3.1 is largely inactivated (14, 51). We found a significant increase in residual T-type currents in *CACNA1G* M1531V neurons compared with control neurons (Fig. 3G).

Additionally, we performed extracellular recordings of control, *CACNA1G* M1531V, and *CACNA1G* KO hDiOs to assess spontaneous neuronal activity (Fig. 3H). We detected increased spontaneous firing rates in neurons carrying both *CACNA1G* M1531V and *CACNA1G* KO variants compared to control neurons (Fig. 3I, J). We also observed increased firing rates in heterozygous forms of *CACNA1G* M1531V and *CACNA1G* KO (Supplementary Fig. 6A-F). In initial experiments (52), we detected hypersynchronous burst-like activity in *CACNA1G* M1531V hDiO, but this phenotype was not consistent across a larger cohort of organoids. Taken together, these findings suggest that different pathogenic variants in *CACNA1G* lead to abnormal channel function and result in changes in spontaneous activity patterns of thalamic neurons.

### Altered connectivity in thalamo-cortical assembloids carrying ***CACNA1G*** pathogenic variants

To investigate circuit-level consequences of the *CACNA1G* variants on thalamocortical projections, we generated thalamo-cortical assembloids and recorded spontaneous neuronal activity via extracellular recordings (Fig. 4A). We found increased spontaneous activity of thalamic neurons in *CACNA1G* M1531V and *CACNA1G* KO thalamo-cortical assembloids (Fig. 4B). Interestingly, the hyperactivity was also observed on the cortical side of the assembloids (Fig. 4C). Notably, the firing rates of cortical neurons in unfused hCO remained unaffected by *CACNA1G* genetic manipulations (Supplementary Fig. 6G-I) suggesting that aberrant hyperactivity may originate on the thalamic side. Moreover, we examined the correlation between thalamic and cortical activity using scaled correlation analysis to investigate whether activity synchrony is affected by *CACNA1G* variants. We found that the averaged correlation values per unit of a-hDiO to a-hCO pairs was higher in *CACNA1G* M1531V, but not in *CACNA1G* KO assembloids (Supplementary Fig. 7A). These results imply that *CACNA1G* M1531V leads to correlated hyperactivity in thalamocortical assembloids, while *CACNA1G* loss-of-function results in uncorrelated hyperactivity.

**Figure 4.**
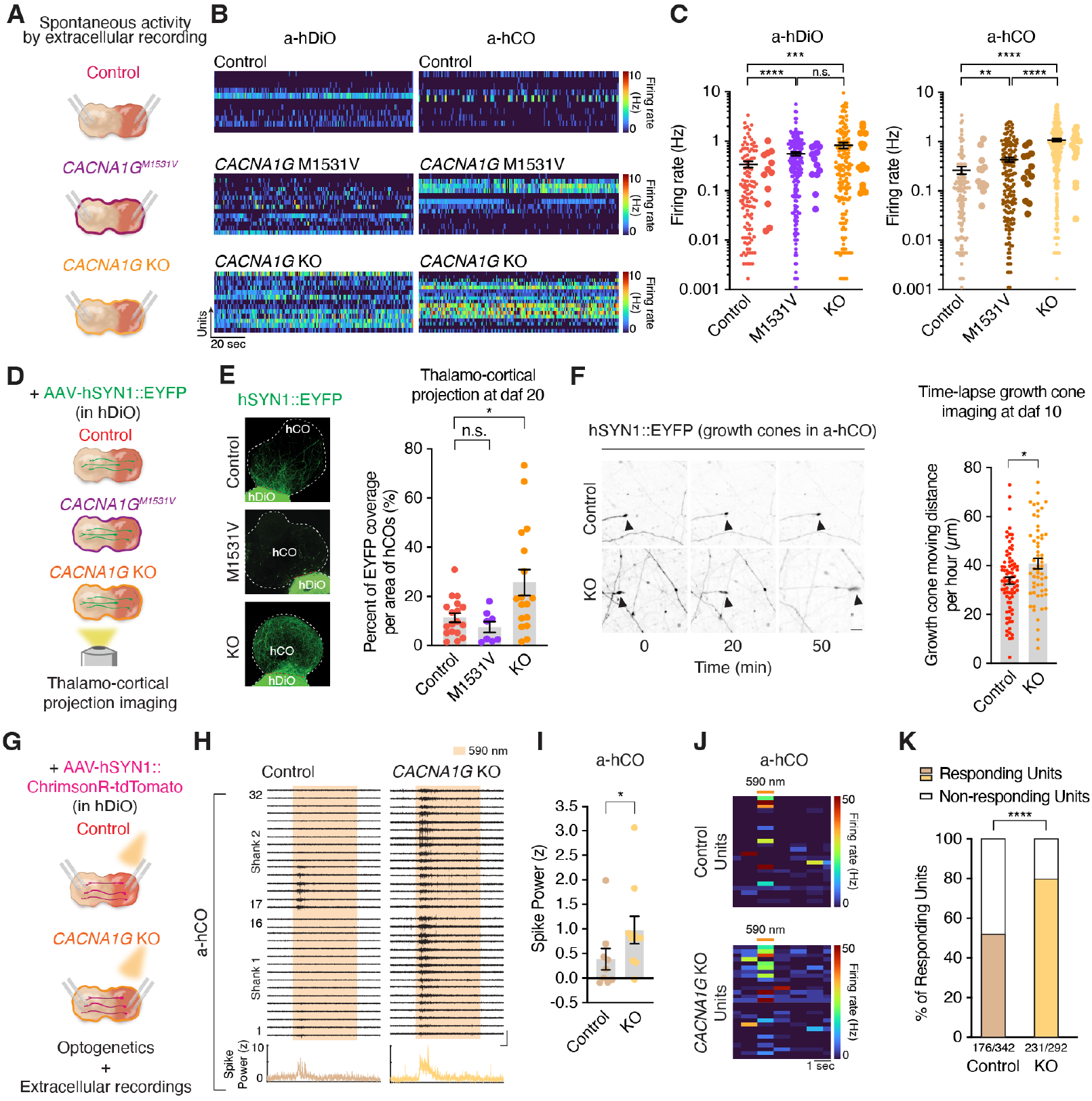
Altered neuronal activity and connectivity in thalamo-cortical assembloids with *CACNA1G* pathogenic variants. (**A**) Schematic illustrating extracellular recording from control, *CACNA1G* M1531V and *CACNA1G* KO hCO–hDiO assembloids. (**B**) Representative heatmaps of spontaneous activity in control (top), *CACNA1G* M1531V (middle), and *CACNA1G* KO (bottom) hCO–hDiO. (**C**) Firing rates of sorted single units from control, *CACNA1G* M1531V, and *CACNA1G* KO a-hCOs (n = 129 units from 12 a-hCOs, 9 differentiations for control, n = 169 units from 13 a-hCOs, 5 differentiations for *CACNA1G* M1531V and n = 308 units from 16 a-hCOs, 7 differentiations for *CACNA1G* KO; Kruskal-Wallis test, **P = 0.0054, ****P < 0.0001), and a-hDiOs (n = 116 units from 12 a-hDiOs, 9 differentiations for control, n = 208 units from 13 a-hDiOs, 5 differentiations for *CACNA1G* M1531V and n = 163 units from 16 a-hDiOs, 7 differentiations for *CACNA1G* KO; Kruskal-Wallis test, ***P = 0.0006, ****P < 0.0001) at daf 35–136. (**D**) Schematic illustrating quantification of thalamocortical projections in hCO–hDiO assembloids. (**E**) Representative images showing thalamocortical projections in hCO–hDiO at daf 20 (left) and their quantification (*n* = 18 assembloids for control, n = 8 assembloids for *CACNA1G* M1531V and n = 17 assembloids for *CACNA1G* KO; Kruskal-Wallis test; n.s., not significant, *\*P* = 0.0451). (**F**) Representative images of time lapse growth cone dynamic (left) and quantification at daf 10 (right) (n = 82 growth cones from 9 assembloids for control, n = 57 growth cones from 6 assembloids for *CACNA1G* KO; Mann-Whitney test, \**P* = 0.0129). Scale bar: 10 μm. (**G**) Schematic illustrating optogenetic stimulation and extracellular recordings in hCO–hDiO. **(H)** Representative extracellular recording traces from control and *CACNA1G* KO assembloids highlighting the response of a-hCO by a-hDiO optogenetic stimulation (top) and z-scored power of spike band frequency (bottom). Vertical scale bar: 100 μV, horizontal scale bar: 100 ms. (**I**) Quantification of z-scored power of spike band frequency in a-hCO during light stimulation at daf 35–85 (*n* = 9 assembloids for control and *n* = 10 assembloids for *CACNA1G* KO; Mann-Whitney test, \**P* = 0.0435). (**J**) Firing rate heatmap of sorted units from representative control and *CACNA1G* KO a-hCOs. (**K**) Percentage of responding units of a-hCO after optogenetic stimulation of the counterpart a-hDiO (n = 342 units from 9 assembloids for control and n = 292 units from 10 assembloids for *CACNA1G* KO; Chi-square test, \*\*\*\**P* < 0.0001). Data is shown as mean ± s.e.m. Small dots indicate units and large dots indicate assembloids in C. Dots indicate assembloids in E, I and growth cones in F.

This prompted us to investigate if additional defects, such as abnormal projections, might contribute to this functional defect. Therefore, we next probed anatomic and functional thalamocortical connectivity in assembloids derived from CACNA1G variant lines. Interestingly, confocal live imaging of thalamocortical projections in the assembloids (Fig. 4D), revealed an increase of projections in *CACNA1G* KO assembloids, but not in *CACNA1G* M1531V assembloids (Fig. 4E). To verify if this was related to abnormal axonal growth, we performed live imaging of growth cone dynamic in *CACNA1G* KO assembloids. We found that *CACNA1G* KO growth cones moved longer distances over time in lived-imaged assembloids (Fig. 4F), but this was not present in *CACNA1G* M1531V assembloids, which is consistent with the projection data (Supplementary Fig 7B). To investigate whether the effects of *CACNA1G* KO on thalamocortical projection formation are autonomous, we generated mosaic assembloids by fusing control hCO with CACNA1G KO hDiO (hCO^Ctrl^–hDiO^*CACNA1G*-KO^) (Supplementary Fig 7C). We found a tendency for increased projections in these combined assembloids (Supplementary Fig 7C) and increased growth cone dynamics (Supplementary Fig 7D). These results suggest that aberrant projections may result from changes in thalamic neurons. To examine whether increased thalamocortical projections in *CACNA1G* KO assembloids contribute to changes in functional connectivity, we recorded neuronal responses in a-hCO following optogenetic stimulation of ChrimsonR-tdTomato expressing a-hDiO (Fig. 4G). We detected an increase in neuronal activity in *CACNA1G* KO a-hCO. The power in the high-frequency band (spike power) was higher in *CACNA1G* KO a-hCO, but not in a-hDiO, following light stimulation (Fig. 4H, I, Supplementary Fig. 7E, F). Furthermore, the proportion of sorted single units that responded to light was higher in *CACNA1G* KO than control assembloids (Fig. 4J, K). Altogether, experiments in thalamo-cortical assembloids illustrate a range of cellular- and circuit-level defects associated with *CACNA1G* channelopathies: a gain-of-function variant caused hyperactivity with correlated activity pattern in the thalamocortical pathway, while the loss of *CACNA1G* resulted in abnormal projections and activity of neurons in the thalamocortical pathway.

## Discussion

To elucidate the pathophysiology of human developmental channelopathies and to develop effective therapeutics, it is necessary to investigate the degree of ion channel dysfunction (i.e., gain- versus loss-of-function), cell-type specific expression and susceptibility, as well as circuit level consequences and compensation. In this study, we probed the cellular- and circuit-level consequences of *CACNA1G* pathogenic variants associated with different clinical presentations in a new human model of thalamocortical connectivity. In-frame deletion and protein-truncating variants in *CACNA1G* have been associated with intellectual disability and schizophrenia (12, 13). We found that deletion of *CACNA1G* resulted in accelerated growth cone movement of thalamic neurons and increased projections in thalamo-cortical assembloids. Surprisingly, loss of this thalamus-enriched calcium channel also resulted in higher hDiO activity. Ca_V_3.1 plays an important role not only in burst firing but also in tonic repetitive firing in thalamic neurons (53). Loss of *CACNA1G* and T-type calcium currents may elicit compensatory mechanisms, engaging other voltage-gated ion channels and could lead to the increased firing rate we observed in this study. Taken together, these defects suggest abnormal thalamocortical connectivity, which is consistent with functional imaging reports (54-57). In contrast, the M1531V point mutation associated with neonatal-onset epilepsy resulted in the slow decay of T-type currents as well as hyperactivity in thalamic neurons, which were simultaneously present in cortical neurons in assembloids. This aligns with pathophysiological models of absence seizures that arise from T-type calcium channel-dependent burst firing of reciprocally connected neurons in the thalamus and neocortex (7).

There are several advantages to our human cellular model. It enables a multi-level assessment of genetic disease variants, including cell type-specific effects in regionalized organoids to inter-regional interactions in assembloids and the consequences (and potential compensation) in thalamocortical networks. As we have shown before, assembloid models are modular, allowing disentanglement of cell autonomous and non-autonomous effects. Our experimental system also allows simultaneous probing of interconnected neuronal populations.

Moving forward, the current assembloids system can be further improved. There are key GABAergic populations in both cortical circuits and in the thalamus (i.e., reticular neurons). Future studies could incorporate these cells through additional assembloid components (58). The thalamocortical circuit is part of a larger loop circuit, thus integration with striatal cells and other GABAergic cells of the midbrain could enable analysis of emergent features of these circuits. The development of thalamocortical circuitry is characterized by waves of activity that spread from the thalamus even in the absence of sensory input, which is thought to contribute to maturation and refinement of connections (59, 60). Larger field imaging coupled with integration of sensory input through either assembly with sensory cells or following transplantation may enable the study of wave-like phenomena and how these shape circuit formation (61).

There are several aspects that we would like to point out when planning experiments with thalamic organoids or thalamo-cortical assembloids. First, there are currently several methods to generate thalamic neurons from pluripotent stem cells in both mouse and human (22, 23). Our differentiation protocol modulates SHH (SAG), WNT (CHIR), and BMP (BMP7) signaling to derive thalamic neurons and inhibits Notch signaling with DAPT to promote neuronal maturation. Other approaches use a combination of BMP7, insulin, and PD0325901 without DAPT. These different protocols may generate different cell types and states that may be useful in the future for modeling different aspects of development and disease. Second, implementing a quality control step early in differentiation, such as assessing the TCF7L2^+^ percentage or *TCF7L2* gene expression in organoids, can be beneficial by increasing phenotyping reliability. Third, it may also be useful to inspect and ideally quantify viral expression and/or projection levels before conducting functional assays in hCO-hDiO assembloids to reduce variability. Overall, this in vitro platform has the potential to reveal cell and network level defects associated with genetic and environmental perturbations at early, previously inaccessible but critical stages of thalamocortical crosstalk in human development and, ultimately, to be used for testing of therapeutics.

## Acknowledgements

We thank members of the Pasca laboratory at Stanford University for scientific inputs and the Stanford Wu Tsai Neurosciences Virus Core for production of AAVs. This work was supported by the Welcome Leap 1kD Program (to S.P.P.), Stanford Brain Organogenesis Big Idea Grant from the Wu Tsai Neurosciences Institute (to S.P.P.), the NYSCF Robertson Stem Cell Investigator Award (to S.P.P.), the Kwan Research Fund (to S.P.P.), the Senkut Research Funds (to S.P.P.), the Ludwig Foundation (to S.P.P.), the Chan Zuckerberg Initiative Ben Barres Investigator Award (to S.P.P.), the CZ BioHub Investigator Program (to S.P.P.), a fellowship from the National Research Foundation of Korea (to J.K), TAA Young Investigator Award (to Y.M.), Stanford Medicine Dean’s Fellowships (to Y.M., F.B. and X.M.), and Stanford Maternal & Child Health Research Institute (MCHRI) Postdoctoral Fellowships (to Y.M., O.R., F.B.). This paper was typeset with the bioRxiv word template by @Chrelli: www.github.com/chrelli/bioRxiv-word-template

## Author contributions

J.K., Y.M. and S.P.P. conceived the project and designed experiments. J.K. carried out the differentiation experiments, characterization of hDiO, extracellular recording and disease modeling. Y.M. performed the differentiation experiments, single cell transcriptomics analyses and calcium imaging with optogenetics. M.-Y.L. performed whole cell recordings and analyses. O.R. and J.R.H. contributed to the development of the extracellular recording platform and the analysis pipeline. S.S. and M.H.P. generated and validated *CACNA1G* variant hiPS cell lines. F.B. and X.M. contributed to optimizing the protocol of regionalized organoids. M.V.T. performed the differentiation experiments and characterization of regionalized organoids. S.D.P analyzed optically-evoked calcium imaging results. J.A. and A.M.P. contributed to primary tissue experiments. J.K., Y.M. and S.P.P. wrote the manuscript with input from all authors.

## Competing interest statement

Stanford University has filed a provisional patent application covering the generation of multi-region assembloids. M.H.P. is on the Board of Directors and holds equity in Graphite Bio. M.H.P. serves on the SAB of Allogene Tx and is an advisor to Versant Ventures.

## Materials and Methods

### Characterization and maintenance of hiPS cells

Human induced pluripotent stem (hiPS) cell lines used in this study were validated using methods described in previous studies (45, 62). Genome-wide SNP genotyping was performed using the Illumina genome-wide GSAMD-24v2-0 SNP microarray at the Children’s Hospital of Philadelphia. Cultures were tested for and maintained Mycoplasma free. A total of 5 control hiPS cell lines derived from fibroblasts collected from 5 healthy subjects were used for experiments. One of the control hiPS cell lines was used to generate *CACNA1G* mutant hiPS cell lines (Supplementary Table 1). Approval for these experiments was obtained from the Stanford IRB panel and informed consent was obtained from all subjects.

### Human primary tissue

Human brain tissue samples were obtained under a protocol approved by the Research Compliance Office at Stanford University.

### Generation of hDiO and hCO from hiPS cells

For neural differentiation, hiPS cells were cultured on vitronectin-coated plates (5 µg/mL, Thermo Fisher Scientific, A14700) in Essential 8 medium (Thermo Fisher Scientific, A1517001). Cells were passaged every 4–5 days with UltraPure^TM^ 0.5 mM EDTA, pH 8.0 (Thermo Fisher Scientific, 15575). For the generation of regionalized neural organoids, hiPS cells were incubated with Accutase® (Innovative Cell Technologies, AT104) at 37°C for 7 min and dissociated into single cells. Optionally, 1–2 days before organoid formation, hiPS cells were exposed to 1% dimethylsulfoxide (DMSO) (Sigma-Aldrich, 472301) in Essential 8 medium. For aggregation into organoids, approximately 3 x 10^6^ single hiPS cells were added per AggreWell-800 well in Essential 8 medium supplemented with the ROCK inhibitor Y27632 (10 µM, Selleckchem, S1049), centrifuged at 100 *g* for 3 min, and then incubated at 37°C in 5% CO_2_. After 24 hours, organoids consisting of approximately 10,000 cells were collected from each microwell by pipetting medium in the well up and down with a cut P1000 pipet tip and transferred into ultra-low attachment plastic dishes (Corning, 3262) in Essential 6 medium (Thermo Fisher Scientific, A1516401) supplemented with the SMAD pathway inhibitors dorsomorphin (2.5 µM, Sigma-Aldrich, P5499) and SB-431542 (10 µM, R&D Systems, 1614). For the first 5 days, Essential 6 medium was changed every day and supplemented with dorsomorphin and SB-431542. To generate hDiO, medium was supplemented with 1 µM CHIR (Selleckchem, S1263) on day 5. On day 6, neural organoids were transferred to neural media containing Neurobasal^TM^-A Medium (Thermo Fisher Scientific, 10888022), B-27^TM^ Supplement, minus vitamin A (Thermo Fisher Scientific, 12587010), GlutaMAX^TM^ Supplement (1:100, Thermo Fisher Scientific, 35050079), Penicillin-Streptomycin (1:100, Thermo Fisher Scientific, 15070063), and supplemented with 1 µM CHIR (Selleckchem, S1263). On day 9 of differentiation, medium was supplemented with 100 nM SAG (day 9–15, Millipore Sigma, 566660-1MG), in addition to the compounds described above. Furthermore, on day 12 of differentiation, organoids were supplemented with 30 ng/mL BMP7 (PeproTech, 120-03P), in addition to the compounds described above. From day 19, to promote differentiation of the neural progenitors into neurons of hDiO, the neural medium was supplemented with brain-derived neurotrophic factor (BDNF; 20ng mL^−1^, PeproTech, 450-02), NT3 (20 ng mL^−1^, PeproTech, 450-03), L-Ascorbic Acid 2-phosphate Trisodium Salt (AA; 200 µM, FUJIFILM Wako Chemical Corporation, 323-44822), N6, 2’-O-Dibutyryladenosine 3’, 5’ -cyclic monophosphate sodium salt (cAMP; 50 µM, Millipore Sigma, D0627), cis-4, 7, 10, 13, 16, 19-Docosahexaenoic acid (DHA; 10 µM, Millipore Sigma, D2534), and 2.5 µM DAPT (only from day 19-25, STEMCELL Technologies, 72082). From day 43, only neural medium containing B-27^TM^ Plus Supplement (Thermo Fisher Scientific, A3582801) was used and changed every 4 days. hCO were generated as previously described (45). From day 22, the neural media was supplemented with BDNF NT3, AA, cAMP, and DHA. From day 46, only neural media containing B-27^TM^ Plus Supplement was used for media changes (every 4 days).

### 3D clearing and staining of a human primary brain sample

To optically clear and stain human primary tissue, the hydrophilic chemical cocktail-based CUBIC protocol was used (63). A dissected primary brain sample was fixed with a 4% PFA/PBS solution at 4°C overnight. The next day, the tissue was washed twice with PBS and incubated in Tissue-Clearing Reagent CUBIC-L (TCI, T3740) at 37°C for 2 days. The sample was washed three times with PBS, then nuclei were stained with RedDot2 (Biotium, #40061, 1:200 dilution) in PBS containing 500 mM NaCl at 37°C overnight. The tissue was washed twice with PBS, and once with a solution containing 10 mM HEPES, 10% TritonX-100, 200 mM NaCl and 0.5% BSA (HEPES-TSB) at 37°C for 2 hours, and then stained with anti-TCF7L2 (rabbit, Cell Signaling Technology, 2569S, 1:500 dilution) antibody in HEPES-TSB solution at 37°C for 2 days. Stained tissue was washed twice with 10% Triton X-100 in PBS and once with HEPES-TSB solution for 2 hours, and then incubated with a donkey anti-rabbit IgG (H&L) highly cross-adsorbed secondary antibody, Alexa Fluor 568 (Thermo Fisher Scientific, A10042, 1:300 dilution) in HEPES-TSB solution at 37°C for 2 days. The sample was washed twice with 10% Triton X-100 in PBS for 30 min and once with PBS for 1 hour. After washing with PBS, the sample was incubated with Tissue-Clearing Reagent CUBIC-R^+^ (TCI, T3741) at room temperature for 2 days for refractive index matching. The CUBIC-cleared tissue was imaged using a 10x objective on a Leica TCS SP8 confocal microscope.

### Cryosection and immunocytochemistry

hCO, hDiO and assembloids were fixed in 4% paraformaldehyde (PFA)/phosphate buffered saline (PBS) overnight at 4°C. They were then washed in PBS and transferred to 30% sucrose/PBS for 2–3 days until the organoids/assembloids sank into the solution. Subsequently, they were rinsed in optimal cutting temperature (OCT) compound (Tissue-Tek OCT Compound 4583, Sakura Finetek) and 30% sucrose/PBS (1:1), embedded, and snap-frozen using dry ice. For immunofluorescence staining, 30 µm-thick sections were cut using a Leica Cryostat (Leica, CM1860). Cryosections were washed with PBS to remove excess OCT from the sections and blocked in 10% Normal Donkey Serum (NDS, Millipore Sigma, S30-100ML), 0.3% Triton X-100 (Millipore Sigma, T9284-100ML), and 0.1% BSA diluted in PBS for 1 hour at room temperature. The sections were then incubated overnight at 4°C with primary antibodies diluted in PBS containing 2% NDS and 0.1% Triton X-100. PBS was used to wash the primary antibodies and the cryosections were incubated with secondary antibodies in PBS with the PBS containing 2% NDS and 0.1% Triton X-100 for 1 hour. The following primary antibodies were used for staining: anti-TCF7L2 (rabbit, Cell Signaling Technology, 2569S, 1:500 dilution), anti-GFP (chicken, GeneTex, GTX13970, 1:1000 dilution), anti-VGLUT2 (mouse, MilliporeSigma, MAB5540, 1:200 dilution), and anti-mCherry (rabbit, GeneTex, GTX128508, 1:1000 dilution). Alexa Fluor dyes (Life Technologies) were used at 1:1,000 dilution and nuclei were visualized with the Hoechst 33258 dye (Life Technologies, H3549, 1:10,000 dilution). Cryosections were mounted for microscopy on glass slides using Aqua-Poly/Mount (Polysciences, 18606), and imaged on a Leica TCS SP8 confocal microscope. Images were processed and analyzed using Fiji (NIH) and IMARIS (Oxford Instruments). hiPS cell cultures on vitronectin-coated glass coverslips were fixed in 4% PFA in PBS for 20 min and then rinsed twice for 5 min with PBS. Coverslips were blocked in 10% NDS, 0.3% Triton X-100, and 0.1% BSA diluted in PBS for 1 hour at room temperature and then incubated overnight at 4°C with a primary antibody, anti-OCT4 (C30A3) (rabbit, Cell Signaling Technology, 2840, 1:200 dilution) diluted in PBS containing 2% NDS and 0.1% Triton X-100. The next day, the solution (including the primary antibody) was washed with PBS three times, and the coverslips were incubated with secondary antibodies including Rhodamin Phalloidin 555 (Thermo Fisher Scientific, #R415, 1:400 dilution) in PBS with 2% NDS and 0.1% Triton X-100 for 1 hour. After PBS washing three times, nuclei were visualized with Hoechst 33258 (Life Technologies, H3549, 1:10,000 dilution), and the coverslips were mounted on glass slides using Aqua-Poly/Mount (Polysciences, 18606), and imaged on a Leica Stellaris 5 confocal microscope.

### Real-time qPCR

Three to five organoids were pooled in the same tube and considered as one sample. mRNA from hCO and hDiO were isolated using the RNeasy Mini kit (Qiagen, 74106) with DNase I, Amplification Grade (Thermo Fisher Scientific, 18068-015). Template cDNA was prepared by reverse transcription using the SuperScript^TM^ III First-Strand Synthesis SuperMix for qRT-PCR (Thermo Fisher Scientific, 11752250). qPCR was performed using the SYBR^TM^ Green PCR Master Mix (Thermo Fisher Scientific, 4312704) on a QuantStudio^TM^ 6 Flex Real-Time PCR System (Thermo Fisher Scientific, 4485689). Primers used in this study are listed in Supplementary Table 2.

### Single cell RNA-seq library preparation and data analysis

Dissociation of hDiO and human tissue was performed as described previously (*64, 65*). For the organoid dissociation, 4–5 randomly selected organoids were pooled to obtain single cell suspension from hDiO and then incubated in 30 U/mL papain enzyme solution (Worthington Biochemical, LS003126) and 0.4% DNase (12,500 U/mL, Worthington Biochemical, LS2007) at 37°C for 45 minutes. After enzymatic dissociation, organoids were washed with a solution including protease inhibitor and gently triturated to achieve a single cell suspension. Cells were resuspended in 0.04% BSA/PBS (Millipore-Sigma, B6917-25MG) and filtered through a 70 µm Flowmi Cell Strainer (Bel-Art, H13680-0070) and the number of cells were counted. To target 7,000 cells (for hDiO) and 10,000 cells (for primary sample) after recovery, approximately 11,600 and 16,000 cells were loaded, respectively, per lane on a Chromium Single Cell 3′chip (Chromium Next GEM Chip G Single Cell Kit, 10x Genomics, PN-1000127) and cDNA libraries were generated with the Chromium Next GEM Single Cell 3' GEM, Library & Gel Bead Kit v3.1 (10x Genomics, PN-1000128), according to the manufacturer’s instructions. Each library was sequenced using the Illumina NovaSeq S4 2 × 150 bp by Admera Health. Quality control, UMI counting of Ensembl genes and aggregation of samples were performed by the ‘count’ and ‘aggr’ functions in Cell Ranger software (version, Cellranger-6.0.1). Further downstream analyses were performed using the R package Seurat. Genes on the X or Y chromosome were removed from the count matrix to avoid biases in clustering due to the sex of the hiPS cell lines. Cells with more than 7,500 or less than 200 detected genes or with mitochondrial content higher than 15% were excluded. Genes that were not expressed in at least three cells were not included in the analysis. Gene expression was normalized using a global-scaling normalization method (normalization method, ‘LogNormalize’; scale factor, 10,000), and the 2,000 most variable genes were selected (selection method, ‘vst’) and scaled (mean = 0 and variance = 1 for each gene) before principal component analysis. The top 10 principal components were used for clustering (resolution of 1.0), using the ‘FindNeighbors’ and ‘FindClusters’ functions, and for visualization with UMAP. Clusters were grouped based on the expression of known marker genes. Unbiased spatial mapping of the whole combined cluster was performed using VoxHunt (version, VoxHunt1.0.0). Briefly, the 50 most variable features from the ISH Allen Brain Atlas data of the E15.5 mouse brain were selected, and similarity maps were calculated. These maps were then plotted in the sagittal views. Comparison to BrainSpan transcriptomic data of micro-dissected human brain tissue at postconceptional weeks (PCW) 15–24 was also performed using VoxHunt with default settings. A circular plot was generated using the R package plot1cell (*66*). The R package scCustomize (v1.1.1) (https://samuel-marsh.github.io/scCustomize) was used to generate the UMAP for integrated samples (‘FeaturePlot_scCustom’ function), and to calculate the percentage of clusters per sample (‘Percent_Expressing’ function). Integration of hDiO and primary tissue samples, as well as control and CACNA1G variants scRNA-seq datasets, was performed using a standard Seurat V3 integration workflow (*67*).

### Viral labeling and live cell imaging

Viral infection of 3D neural organoids was performed as previously described (45). Briefly, neural organoids were transferred into a 1.5 mL Eppendorf tube containing 200 µL of neural media and incubated with the virus overnight at 37°C, 5% CO_2_. Next day, 800 µL of fresh culture media was added. The following day, neural organoids were transferred into fresh culture media in ultra-low attachment plates (Corning, 3471, 3261). For live cell imaging, labeled hCO or hDiO or assembloids were transferred into a well of a Corning^TM^ 96-Well microplate (Corning, 4580) in 150 µL of neural media or on a 20 mm glass coverslip in a 35 mm glass bottom well (Cellvis, D35-20-0-N) and incubated in an environmentally controlled chamber for 15–30 min before imaging on a Leica TCS SP8 or Leica Stellaris 5 confocal microscope. The viruses used in this study were: AAV-DJ-hSYN1::EYFP (Stanford University Neuroscience Gene Vector and Virus Core, GVVC-AAV-16), AAV-DJ-hSYN1::mCherry (Stanford University Neuroscience Gene Vector and Virus Core, GVVC-AAV-17), AAV-DJ-EF1-DIO-tdTomato (Stanford University Neuroscience Gene Vector and Virus Core, GVVC-AAV-169), AAV1-hSYN1::Cre (Addgene, #105553), AAV1-hSYN1::ChrimsonR-tdTomato (68) (Addgene, #59171-AAV1) and AAV-DJ-hSYN1::GCaMP6f (Stanford University Neuroscience Gene Vector and Virus Core, GVVC-AAV-96).

### Generation of thalamo-cortical assembloids

To generate thalamo-cortical assembloids, hDiO and hCO were separately generated from hiPS cells and after day 46, hDiO and hCO were assembled by placing them in close proximity in 1.5 mL Eppendorf tubes, inside an incubator, for 3 days. Medium was gently replaced on day 2. On the third day, assembloids were transferred using a cut P1000 pipette tip in 24-well ultra-low attachment plates in the neural media described above. Media was changed every 4 days.

### Axon projection imaging in thalamo-cortical assembloids

AAV-DJ-hSYN1::EYFP^+^ cells projecting into hCOs from hDiOs or AAV-DJ-hSYN1::mCherry^+^ cells projecting into hDiOs from hCOs were imaged under environmentally controlled conditions in intact thalamo-cortical assembloids using a Leica TCS SP8 or Leica Stellaris 5 confocal microscope with a motorized stage. Assembloids were transferred to a well in a 24-well glass bottom plate (Cellvis, P24-0-N) in culture medium, and incubated in an environmentally controlled chamber for 15–30 min before imaging. Images were taken using a 10x objective to capture the entire a-hCO or a-hDiO side at a depth of 100 µm. a-hDiO or a-hCO-derived projections were quantified using Fiji (ImageJ, version 2.1.0, NIH). ROIs were manually drawn to cover the area on the a-hDiO or a-hCO to be measured in maximal projection confocal stacks. The percentage of fluorescence positive pixels over total area of a-hDiO or a-hCO was calculated in binary images with consistent threshold values across images.

### Viral transsynaptic tracing in thalamo-cortical assembloids

For anterograde tracing in thalamo-cortical assembloids, hCO were infected with AAV-DJ-EF1α::DIO-tdTomato and AAV-DJ-hSYN1::EYFP, while hDiO were infected with AAV1-hSYN1::Cre. For retrograde tracing in thalamo-cortical assembloids, hCO were infected with rabies-ΔG-Cre-EGFP and AAV-DJ-EF1α::CVS-G-WPRE-pGHpA, while hDiO were infected with AAV-DJ-EF1α::DIO-tdTomato. After viral infection, hCO and hDiO were assembled and maintained in culture with medium changes every 4 days. After 15–38 days after assembly, z-stack images from assembloids were captured using a Leica TCS SP8 or Leica Stellaris 5 confocal microscope and analyzed in Fiji (ImageJ, version 2.1.0, NIH).

### Optogenetic stimulation and calcium imaging

Thalamo-cortical assembloids expressing AAV1-hSYN1::ChrimsonR-tdTomato in hDiO and AAV-DJ-hSYN1::GCaMP6f in hCO were placed on a 20 mm coverslip glass (in a 35 mm glass-bottom plate) in neural medium, and imaged using a 10x objective on a Leica TCS SP8 confocal microscope. For optogenetic stimulation, ChrimsonR-tdTomato^+^ cells in hDiO were activated with 590 nm light using an optical fiber-coupled LED (Thorlabs, M590F3). GCaMP6f was imaged at a frame rate of 14.7 frames/sec. Imaging experiments were 1500 frame-long and one 590 nm of LED (71 ms) was applied every 150 frames. The pulse was generated by a Cyclops LED connected to the Leica TCS SP8. Results were analyzed with Fiji (NIH) and MATLAB (version R2022a, MathWorks). After ROI registration, raw time series were transformed to relative changes in fluorescence: ΔF/F_(t)_ = (F_(t)_-F_0_)/F_0_. To verify whether the optically-evoked responses were time-locked to the LED stimulation, ΔF/F responses were compared to ΔF/F values at randomly selected time points. For time-locked ΔF/F, the amplitudes of ΔF/F from each cell were calculated as the maximum ΔF/F value within a 20–30 frames window (1,360 to 2,040 ms) following LED stimulation (minus the mean of the baseline 1 sec before the stimulation). The median was used to exclude shape-dependent artifacts.

### Extracellular recording and analyses

Neural organoids or assembloids were embedded into 3% low-melting gel agarose (IBI Scientific, IB70056) and transferred into artificial cerebrospinal fluid (aCSF) containing 124 mM NaCl, 3 mM KCl, 1.25 mM NaH_2_PO_4_, 1.2 mM MgSO_4_, 1.5 mM CaCl_2_, 26 mM NaHCO_3_ and 10 mM D-(+)-glucose with addition of GlutaMAX (Gibco, 35050061). Embedded neural organoids or assembloids were placed on a Brain Slice Chamber-2 (Scientific Systems Design Inc, S-BSC2) and perfused with aCSF (bubbled with 95% O_2_ and 5% CO_2_) at 37°C. Temperature was controlled and retained at 37°C by connecting to a Proportional Temperature Controller PTC03 (AutoMate Scientific, S-PTC03). Acute 32 channel P-1 probe with 2 shanks (Cambridge Neuro-Tech, ASSY-37 P-1) were connected to an Acute probe adaptor; 32 channels Samtec to Omnetics (Cambridge NeuroTech, ADPT A32-Om32). Signals were acquired using the Intan 1024ch recording controller (Intan Technologies) at 30000 Hz. For optogenetic stimulation, the pulses were generated by the pulse stimulator (Grass Instrument, S48) connected to a Cyclops LED driver and a fiber cable (Thorlabs, M126L01). A 20 sec recording protocol (a 19 sec-long light-off phase followed by a 1 sec-long constant 590 nm illumination) was repeated in 9 trials. Raw recording data (in .rhd format) were opened using the Intan Technologies code and processed in Python. Single unit sorting was performed using Kilosort 4 (69) (version 4.03) with the Spikeinterface package (70) (version 0.7.6). Samples showing large noise or aberrant signals distorting the single-unit sorting process were excluded from further analysis. For spontaneous firing rate analysis, the total number of spikes was divided by the recording duration. To calculate firing rates for optogenetics experiments, the number of spikes during light-on or light-off phases were divided by the duration of light for each phase. Units were considered ‘responding’ if the mean firing rates during the light-on phase was significantly (by paired t-test) increased over the light-off phase over 9 trials. To measure spike power, the continuous wavelet transform analysis (CWT) function in MATLAB (version R2022a, MathWorks) was applied to median signal values obtained from 32 channels; this produced time-frequency decomposition and averaged absolute values of frequency band for spike activity (500 ∼ 2000 Hz), which are referred to as spike power, were z-scored. The spike power of the first 250 ms of each light stimulation was averaged over 9 trials (per assembloid).

### Patch-clamp recordings

Patch-clamp recordings were performed as described previously (*71*). Briefly, hDiOs were infected with AAV-DJ-hSYN1::EYFP and placed on top of cell culture inserts (0.4 mm pore size; Corning, 353090) that were positioned in 6-well plates to form flattened organoids. Recordings were typically made on day 128–135. EYFP^+^ neurons were identified with an upright slice scope microscope (Scientifica) equipped with an Infinity2 CCD camera and Infinity Capture software (Teledyne Lumenera). Recordings were performed with borosilicate glass electrodes with a resistance of 7–10 MΩ. BrainPhys neuronal media (STEMCELL Technologies, 05790) was used as an external solution. The internal solution contained 127 mM K-gluconate, 8 mM NaCl, 4 mM MgATP, 0.3 mM Na_2_GTP, 10 mM HEPES and 0.6 mM EGTA, pH adjusted to 7.2 with KOH (290 mOsm). For hyperpolarization– induced rebound spikes, cells were held at –50 mV and hyperpolarizing current steps (1 sec) with an increment of 10 pA from –100 pA to 0 pA were tested. For barium current recordings, cells were recorded in the presence of TTX (0.5 µM) to block sodium currents, and were held at –70 mV in voltage-clamp and voltage steps (1 sec, from –90 mV to +40 mV) were given with an increment of 10 mV. The external solution contained 100 mM NaCl, 3 mM KCl, 2 mM MgCl_2_, 4 mM BaCl_2_, 25 mM TEA-Cl, 4 mM 4-AP, 10 mM HEPES, 20 mM glucose, pH 7.4 with NaOH, 300 mOsm. The internal solution contained 110 mM CsMethylSO_3_, 30 mM TEA-Cl, 10 mM EGTA, 4 mM MgATP, 0.3 mM Na_2_GTP, 10 mM HEPES, 5 mM QX314-Cl, pH 7.2 with CsOH, 290 mOsm. Leak subtraction was implemented in MultiClamp 700B software. *I-V* curves were fitted in Origin (OriginPro 2021b, OriginLab) with a Boltzmann exponential function: *I* = *G*_*max*_ * (*V* – *E*_*Ba*_) / {1 + exp[(*V*_*0*.*5*_ – *V*) / *K*]}, where *G*_*max*_ is the maximal conductance of the calcium channels, *E*_*Ba*_ is the reversal potential of barium estimated by the curve-fitting program, *V*_*0*.*5*_ is the potential for half-maximal steady-state activation of the barium current, and *K* is a voltage-dependent slope factor. To compare barium current kinetics in control and the CACNA1G M1531V mutant, cells were held at – 70 mV, and voltage steps (with a 10–mV increment) were applied (from –90 mV to +20 mV). Fast T-type currents were measured at –30 mV. Cells showing T-type currents at –30 mV were analyzed. We assume that the fast currents at earlier steps were indicative of Cav3.1 activation with minimal interference from other voltage-gated calcium channels (Cav1, Cav2.1, Cav2.2 or Cav2.3), although some activation cannot be excluded. Data were acquired with a MultiClamp 700B Amplifier (Molecular Devices) and a Digidata 1550B Digitizer (Molecular Devices), low-pass filtered at 2 kHz, digitized at 20 kHz and analyzed with pCLAMP (version 10.6, Molecular Devices). Cells were given a –10 mV hyperpolarization (100 ms) every 10 sec to monitor input resistance and access resistance. Cells were excluded for analysis if either value changed by > 30%. The liquid junction potential was not corrected.

### Genome editing of hiPS cells to generate *CACNA1G* variant hiPS cell lines

For generating *CACNA1G* knockout cells, hiPS cells were genome edited with dual sgRNA:Cas9 RNP complexes to create a frameshift deletion in exon 1 of the *CACNA1G* gene. For this, 24 hours prior to gene editing, hiPS cells were treated with 10 µM ROCK inhibitor (Y-27632). hiPS cells (at 70– 80% confluence) were detached from the culture plate and dissociated into single cells by following incubation with Accutase (Innovative Cell Technologies) for 5 min at 37°C. ROCK inhibitor-supplemented mTeSR1 media was added to neutralize Accutase. Concurrent with the cell dissociation process, the Cas9 RNP complex was prepared. The RNP complex for each of the two sgRNA was formed separately by combining 5 µg of HiFi Cas9 (Aldevron) and 1.75 µg of sgRNA (Synthego) (molar ratio of 1:2.5 (Cas9: sgRNA)) for 10 min at room temperature. The two RNP complexes were then combined and diluted in 20 µL of P3 Primary Cell Nucleofector Solution (Lonza). To electroporate the Cas9 RNP complexes into hiPS cells, 500,000 cells were mixed with the nucleofection solution containing the RNPs and were electroporated in one well of a 16-well Nucleocuvette Strip using the program CA137 in 4D Nucleofector system (Lonza). Following electroporation, cells were plated into two wells of a Matrigel-coated 24-well plate containing 500 µL of mTeSR1 media supplemented with 10 µM Y-27632. The next day, the cells were switched to fresh 10 µM Y-27632 supplemented mTeSR1 medium. After 48 hours of electroporation, cells were cultured in mTeSR1 medium without Y-27632. To isolate single cell clones, edited hiPS cells were detached and dissociated into single cells using Accutase. After dissociation, Accutase was neutralized with mTeSR1 medium and cells were counted. Single cells were then plated at a density of 250 cells per well of a 6-well plate in mTeSR1 medium supplemented with 1X CloneR solution (Stem Cell 28 Technologies) and incubated for 48 hours. At day 2, cells were switched to fresh mTeSR1 with 1X CloneR and incubated for 2 more days. From day 4, cells were cultured in mTeSR1 medium without CloneR. At day 7–10, single cell colonies were picked by scraping and were then plated in one well of a 48-well plate in mTeSR1 medium with 10 µM Y-27632. From the next day, single cell clones were maintained in mTeSR1 medium. The genotype of edited clones was determined by PCR with the primer set FW, 5’-GAAGCGAAGAAGCCGGAACAA-3’; RV, 5’-CCCGGGTTACCGACGAGA-3’, and Sanger sequencing (Mclab or Genewiz) was performed with the following primers: FW, 5’-TAGAGCCCAC-CAGATGTGCC-3’, RV, 5’-GGGCTCACTTCGACTCACC-3’. ICE analysis (https://ice.synthego.com/) was performed on sequencing chromatograms to determine the genotype. Synthetic sgRNA (with 2’-O-methyl 3’- phosphorothioate modified nucleotides at the three terminal positions at both the 5’ and 3’ ends) were purchased from Synthego. The genomic target sequences of the sgRNAs were G2: 5’-GATATGGGTTACAGACCGTG-3’ and G3: 5’-TTGAGCCGCATGAAGCTCCG-3’. sgRNAs with high specificity scores were selected based on the COSMID tool (https://crispr.bme.gatech.edu). For generating the *CACNA1G* M1531V isogenic hiPS cells, AAV6/Cas9 RNP genome editing was performed, as described previously (*50*). Cas9:sgRNA RNP complex was nucleofected into hiPS cells as above. For this gene editing, an sgRNA with a target site close to the genomic region encoding for M1531 was used. After nucleofection, the cells were plated in 10 µM Y-27632-supplemented mTeSR1 medium with a AAV6 homology directed repair donor template at an MOI of 5K, and incubated at 37°C for 24 hours. Cells were then switched to fresh mTeSR1 medium with 10 µM Y-27632. Next day forward, cells were maintained in mTeSR1 medium. Genome edited hiPS cells were subjected to single cell cloning as described above. The single cell clones were genotyped using PCR, Sanger sequencing and ICE analysis. The primer set used for PCR was FW: 5’-CATGGGGTCTCTAGCCGTC-3’ and RV: 5’-GAAAGACTAC-CGACCAGCCA-3’ and the Sanger sequencing primer used was, 5’-CTGTGGGCAGCTCCTAAGACA-3’. ICE analysis was done on the sequencing chromatograms to determine the frequency of control, INDEL and HDR in the single cell clones. The genomic target sequence of the sgRNA used was: 5’-CATCTCGTTCCTGCTCATTG-3’. AAV6 donor template was designed and constructed with 1.2 kb homology arms, incorporating the intended M1531V variant and silent variants on the sgRNA target site to avoid recutting post-HDR. The donor template sequence was cloned into a pAAV-MCS backbone using NEB hifi DNA assembly kit. AAV6 was packaged in 293T cells, purified using the AAVpro purification kit (Takara), and the titer was determined using ddPCR, as described previously (*50*).

## Statistics

Data are presented as mean ± s.e.m. unless described otherwise. Raw data were tested for normality of distribution and statistical analyses were performed by paired and unpaired *t*-test (two-tailed), Mann-Whitney test, Wilcoxon test, one-way *ANOVA*, or Kruskal-Wallis test with multiple comparison tests depending on the dataset. Sample sizes were estimated empirically. GraphPad Prism Version 9.3.1 or MATLAB (version R2022a, MathWorks) were used for statistical analyses.

## Code availability

The codes used for calcium imaging and electrophysiology analyses in this study are available on request from the corresponding author.

---

**Supplementary Figure 1.**
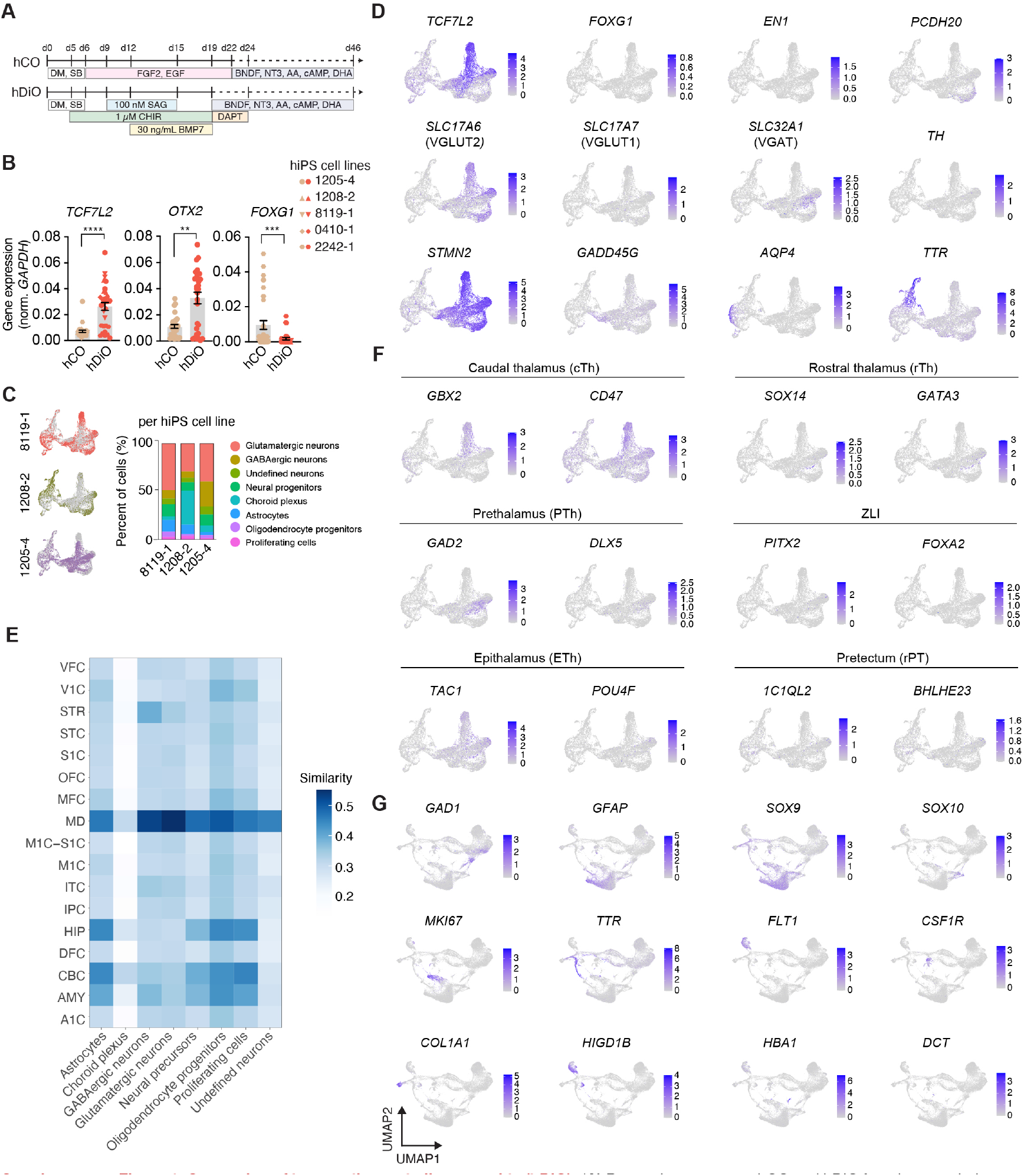
Generation of human diencephalic organoids (hDiO). (**A)** Protocols to generate hCO and hDiO from human pluripotent stem cells. (**B**) Normalized expression of *TCF7L2, OTX2* and *FOXG1* from hCO and hDiO at day 18–19 of differentiation as measured by qPCR (*n* = 33–37 samples from 10 differentiation of 5 hiPS cell lines; Mann-Whitney test, \*\*\**P* = 0.0006 for *OTX2*, \*\*\**P* = 0.0001 for *FOXG1*, \*\*\*\**P* < 0.0001). (**C**) UMAP of hDiO cells by hiPS cell line and distribution of cell clusters. (**D**) UMAP showing cell type markers. (**E**) VoxHunt analysis per cluster. (**F**) UMAP indicating regional thalamic markers. (**G**) *In vitro* and *in vivo* counterpart comparison of scRNAseq data. UMAP including hDiO and GW18–22 primary thalamus. Data is shown as mean ± s.e.m. Dots indicate samples in B. Each shape represents a hiPS cell line: Circle, 1205-4; Triangle, 1208-2; Inverted triangle: 8119-1; Rhombus, 0410-1; Hexagon, 2242-1 in B. Abbreviations: VFC; ventrolateral frontal cortex, V1C; primary visual cortex, STR; striatum, STC; Posterior superior temporal cortex, S1C; primary somatosensory cortex, OFC; orbital frontal cortex, MFC; medial frontal cortex, MD; mediodorsal nucleus of thalamus, M1C; primary motor cortex, ITC; inferior temporal cortex, IPC; posterior inferior parietal cortex, HIP; hippocampus, DFC; dorsolateral prefrontal cortex, CBC; cerebellar cortex, AMY; amygdala, A1C; primary auditory temporal cortex.

**Supplementary Figure 2.**
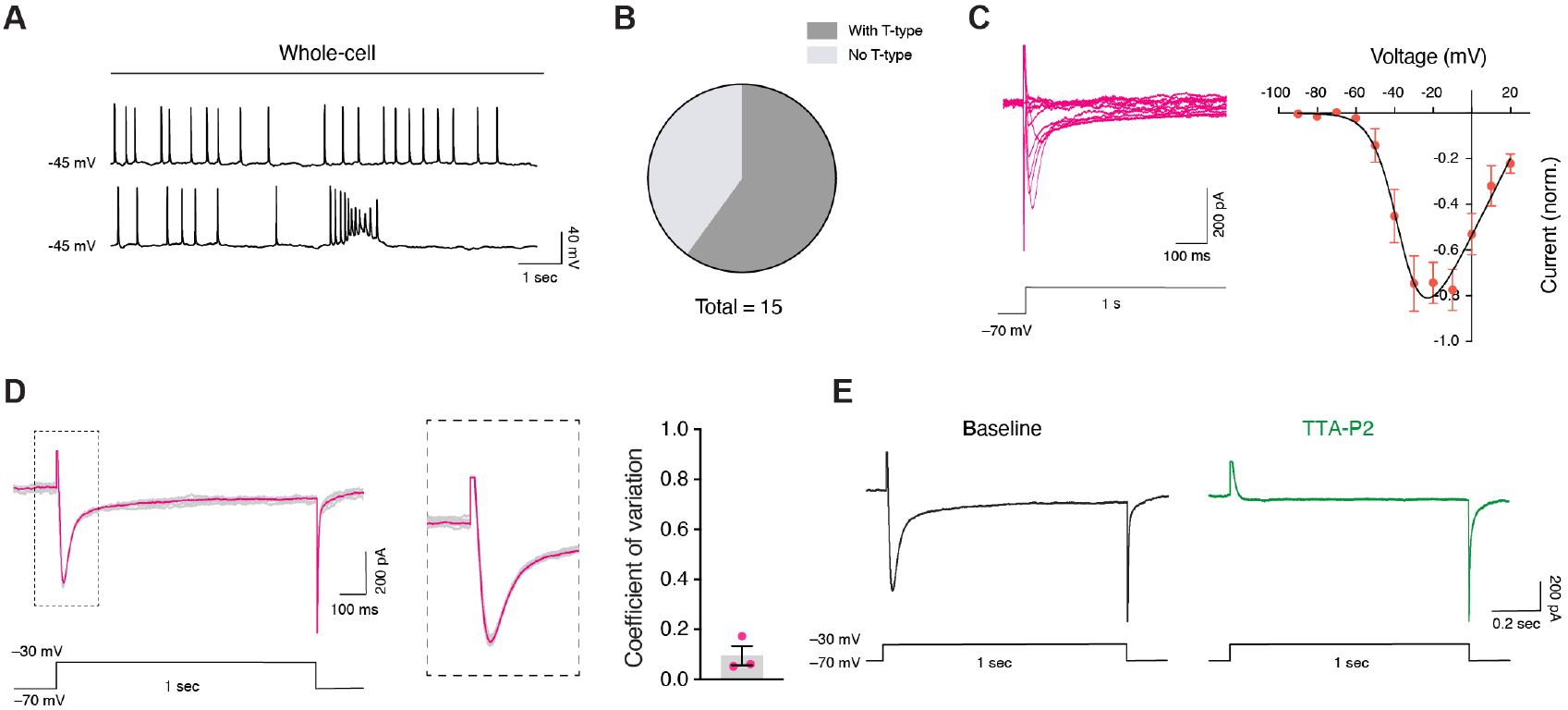
Electrophysiological characterization of hDiO neurons. (**A**) Representative traces showing spontaneous firing of EYFP-labeled hDiO neurons in whole-cell recordings (current clamp). (**B**) Summary of percentage of control hDiO neurons with or without fast T-type barium currents. (**C**) Representative examples and *I–V* curves (barium currents) of control hDiO neurons (held at –70 mV, voltage steps from –90 mV to +20 mV with an increment of 10 mV), n = 7 neurons. (**D**) Representative barium current traces (left) and zoomed-in traces (right). Traces for individual trials and the average are shown in grey and pink respectively. (**E**) Representative averaged barium current traces for depolarization at – 30 mV at baseline (left) and after TTA-P2 treatment (right). The baseline trace is the same recording from the averaged trace in (**D**).

**Supplementary Figure 3.**
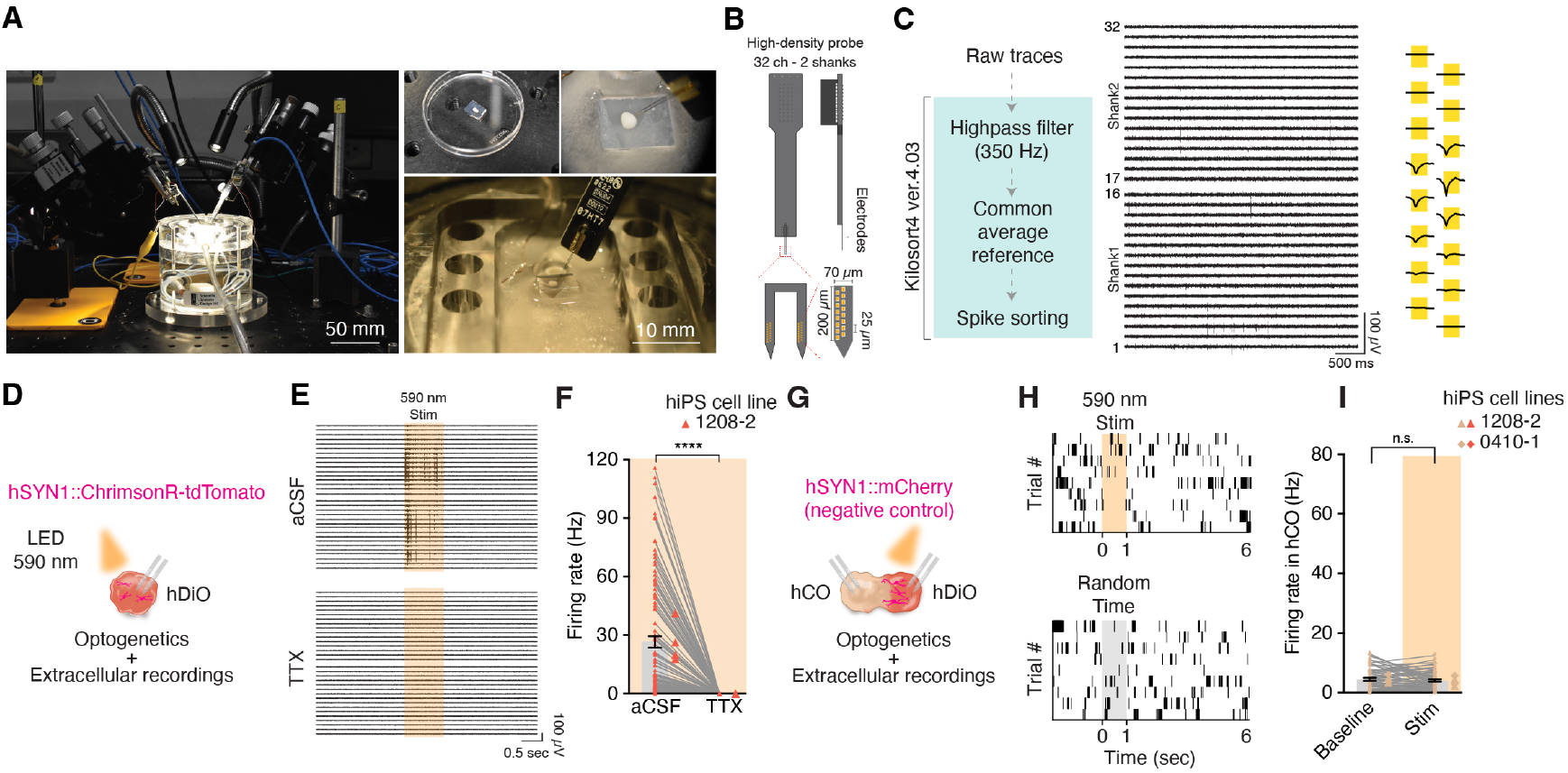
Optogenetics and extracellular recording of thalamo-cortical assembloids. (**A**) Image of a custom extracellular recording platform for organoids and assembloids (left). Organoids are embedded into 3% agarose (top, left and right) and then are placed on the Brain Slice Chamber (bottom, right). (**B**) Design of high-density probe that includes 32 channels in 2 shanks. (**C**) Process for single unit sorting. (**D**) Schematic illustration extracellular recordings combined with optogenetic stimulation of hDiO expressing ChrimsonR-tdTomato. (**E**) Representative trace showing optogenetic stimulation in artificial CSF (top) and TTX treatment (bottom). (**F**) quantification of firing rates in hDiO, before and after TTX treatment (*n* = 106 units from 4 hDiOs, 2 differentiations, 2 hiPS cell lines; Wilcoxon test, \*\*\*\**P* < 0.0001). (**G**) Schematic showing a control experiment with extracellular recordings of a-hCO following light application of to a-hDiO not expressing an opsin in thalamo-cortical assembloids (**H**) Representative raster plot aligned with light stimulation (top) and randomly selected time point (bottom). (**I**) Quantification of firing rates in a-hCO during light on and off phases (*n* = 61 units from 4 assembloids, 1 differentiation of 2 hiPS cell line; Wilcoxon test, n.s., not significant). Data is shown as mean ± s.e.m. Small dots indicate units and large dots indicate organoids or assembloids in F, I. Each shape represents a hiPS cell line: Triangle, 1208-2; Rhombus, 0410-1 in F, I.

**Supplementary Figure 4.**
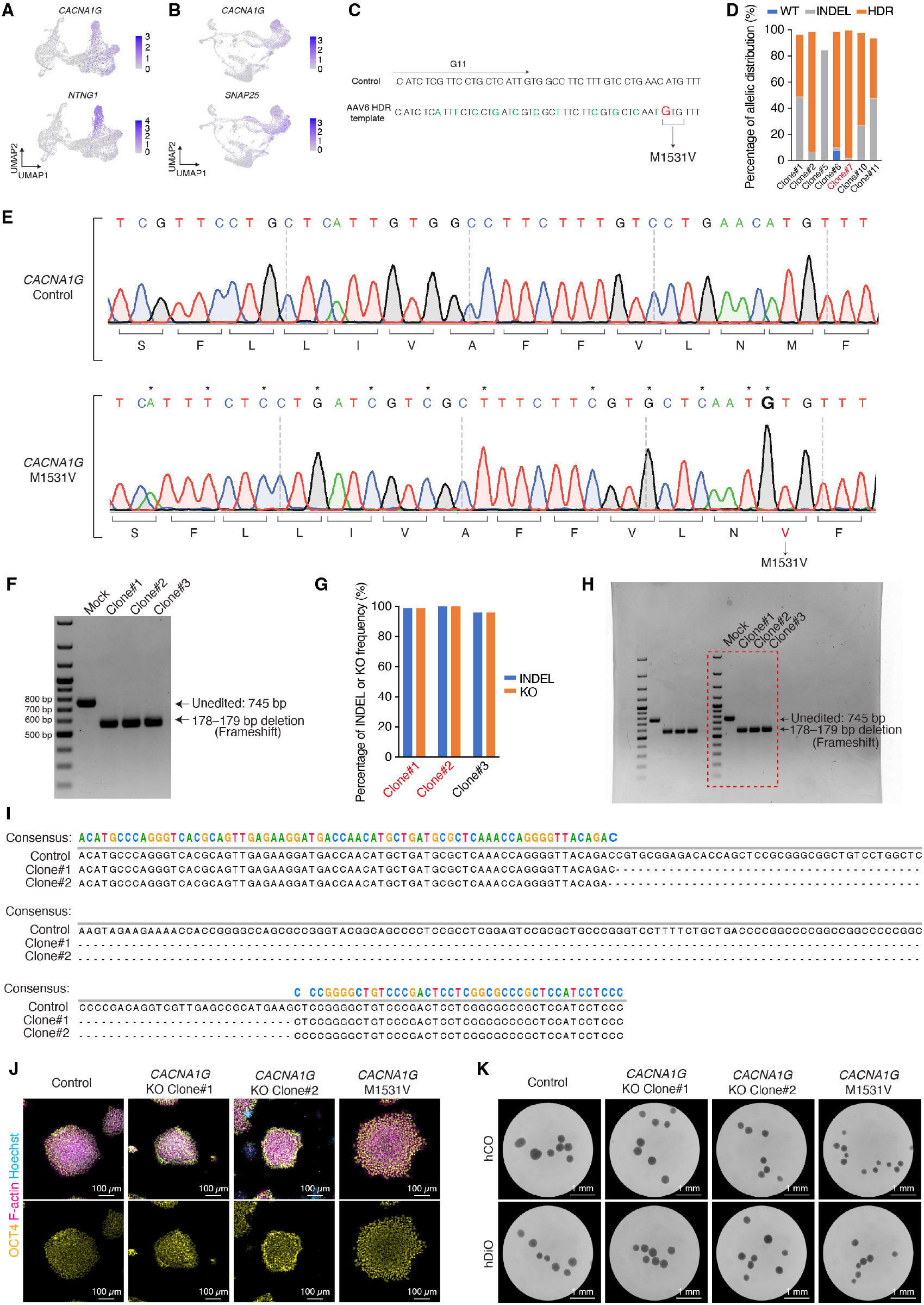
CRISPR/Cas9 based generation of hiPS cell lines carrying *CACNA1G* variants. (**A**) UMAP showing expression of *CACNA1G* (top) and *NTNG1* (bottom) in hDiO at day 100 of differentiation. (**B**) UMAP showing expression of *CACNA1G* (top) and *SNAP25* (bottom) in combined hDiO and human primary thalamus. (**C**) Alignment of the *CACNA1G* control and donor template (AAV6) sequences around the M1531V variant site. AAV6 donor sequence was designed to introduce the desired M1531V variant (in red) and silent variants (in green) to prevent recutting after gene editing. G11 denotes the target site of the sgRNA used for gene editing. (**D**) Allelic distribution of control, INDEL and HDR (M1531V) frequencies based on ICE analysis in the gene edited single cell hiPS cell clones. Clone#7 was used in this study. (**E**) Sanger sequencing results of parental hiPS cells (top) and genome edited *CACNA1G* M1531V hiPS cells (bottom). (**F**) PCR on gene edited hiPS cell clones amplifying exon 1 of *CACNA1G*. Clones#1, #2 and #3 demonstrated a deletion in exon 1. (**G**) Allelic frequency of INDEL and KO based on ICE analysis on the Sanger sequences of PCR amplicons from **F**. This analysis confirmed homozygous KO due to a frameshift deletion of 178/179-bp in clones #1 and #2, which were used in this study. (**H**) Uncropped gel image of **F**. (**I**) Sanger sequencing results of parental hiPS cells (top) and *CACNA1G* KO hiPS cell lines, clone#1 and #2 (middle and bottom). (**J**) Immunostaining of pluripotency marker OCT4 (yellow), F-actin (magenta) and Hoechst (cyan). Scale bar: 100 µm. (**K**) Regionalized organoids generated from the edited hiPS cells. Scale bar: 1 mm.

**Supplementary Figure 5.**
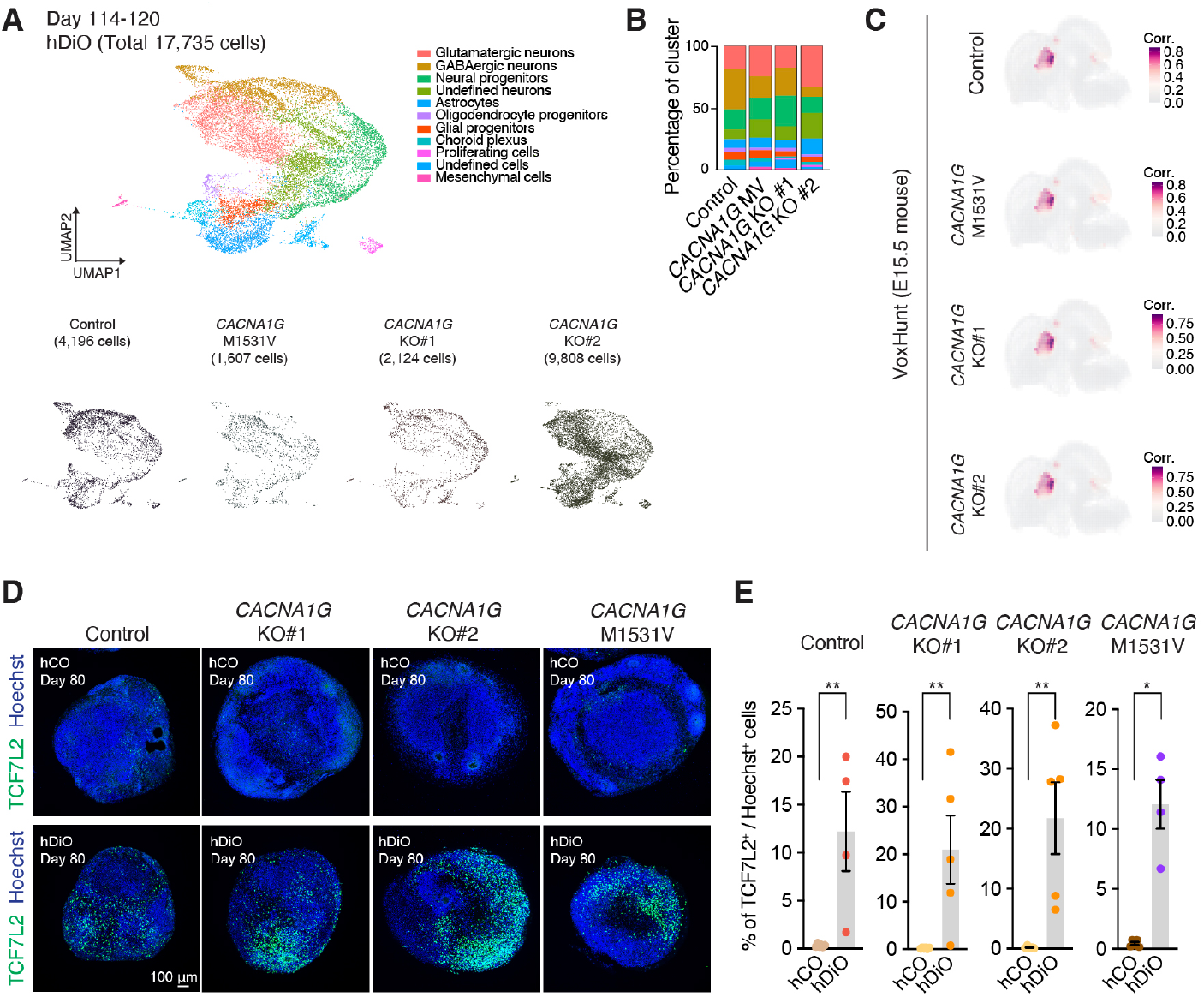
Characterization of control and *CACNA1G* gene variants hDiO. (**A**) UMAP of integrated hDiOs derived from control and CACNA1G gene variants hiPS cell lines. (**B**) Percentage distribution of cell clusters in each sample. (**C**) VoxHunt spatial brain mapping of hDiOs onto the Allen Brain Institute E15.5 mouse brain data. (**D** and **E**) Immunostaining for TCF7L2 (green), and Hoechst (blue) in hCO and hDiO at day 85, and quantification of TCF7L2^+^ cells (n = 5–6 for hCO and n = 3–5 for hDiO from 2 differentiation; Mann-Whitney test, \*\**P* = 0.0095 for control, \*\**P* = 0.0043 for KO#1, \*\**P* = 0.0079 for KO#2, \*\**P* = 0.00159 for M1531V). Scale bar: 100 μm.

**Supplementary Figure 6.**
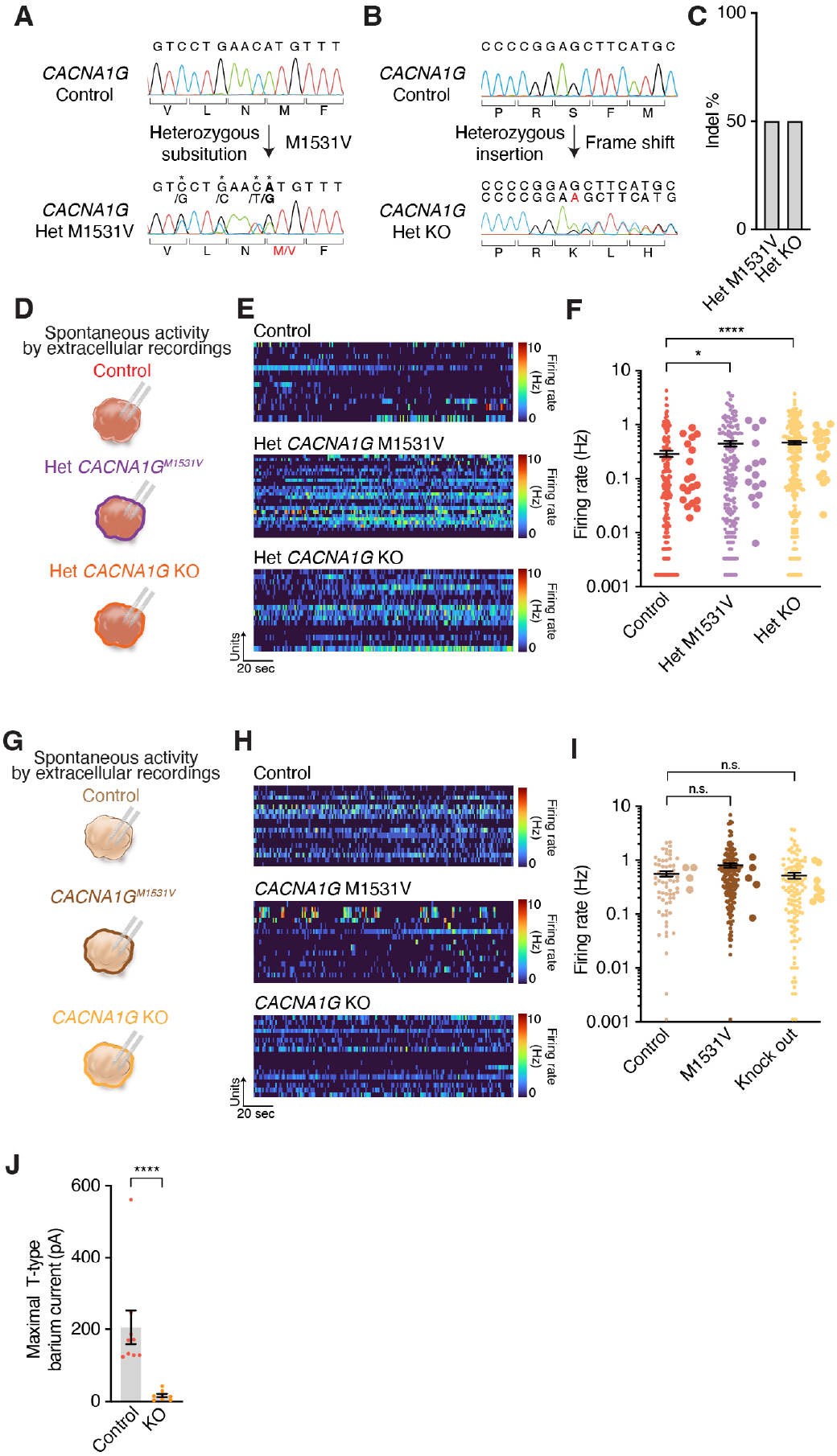
Extracellular recordings from hCOs with *CACNA1G* variants and hDiOs with heterozygous *CACNA1G* variants. (**A**) Sanger sequencing results of control hiPS cells (top) and gene edited heterozygous *CACNA1G* M1531V hiPS cells (bottom). (**B**) Sanger sequencing results of control hiPS cells (top) and gene edited heterozygous *CACNA1G* KO hiPS cells (bottom). (**C**) Allelic frequency of INDEL based on ICE analysis on the Sanger sequences of PCR amplicons from **A** and **B**. (**D**) Schematic illustrating the experimental design for probing spontaneous activity via extracellular recording from control, heterozygous *CACNA1G* M1531V, and heterozygous *CACNA1G* KO hDiOs. (**E**) Representative heatmaps of spontaneous extracellular recording from control (top), heterozygous *CACNA1G* M1531V (middle) and heterozygous *CACNA1G* KO (bottom) hDiOs. (**F**) Firing rates of sorted single units from control, heterozygous *CACNA1G* M1531V, and heterozygous *CACNA1G* KO hDiOs at day 99–176 (n = 203 units from 19 hDiOs, 2 differentiations for control, n = 175 units from 15 hDiOs, 2 differentiations for *CACNA1G* M1531V and n = 243 units from 19 hDiOs, 2 differentiations for *CACNA1G* KO; Kruskal-Wallis test, \*\*\*\**P* < 0.0001). (**G**) Schematic illustrating the experimental design for probing spontaneous activity via extracellular recording from control, *CACNA1G* M1531V, and *CACNA1G* KO hCOs. (**H**) Representative heatmaps of spontaneous extracellular recording from control (top), *CACNA1G* M1531V (middle) and *CACNA1G* KO (bottom) hCOs. (**I**) Firing rates of sorted single units from control, *CACNA1G* M1531V, and *CACNA1G* KO hCOs at day 116–122 (n = 64 units from 4 hCOs, 1 differentiation for control, n = 169 units from 5 hCOs, 2 differentiations for *CACNA1G* M1531V and n = 115 units from 8 hCOs, 2 differentiations for *CACNA1G* KO; Kruskal-Wallis test, \*\*\*\**P* < 0.0001). (**J**) Quantification of maximal T-type barium currents induced by depolarization (from –70 mV) from control and *CACNA1G* KO EYFP^+^ neurons in hDiOs (n = 9 neurons for control; n = 8 neurons for *CACNA1G* KO; Mann-Whitney test, \*\*\*\**P* < 0.0001). Data is shown as mean ± s.e.m. Small dots indicate units or cells in F, I, J and large dots indicate organoids in F, I.

**Supplementary Figure 7.**
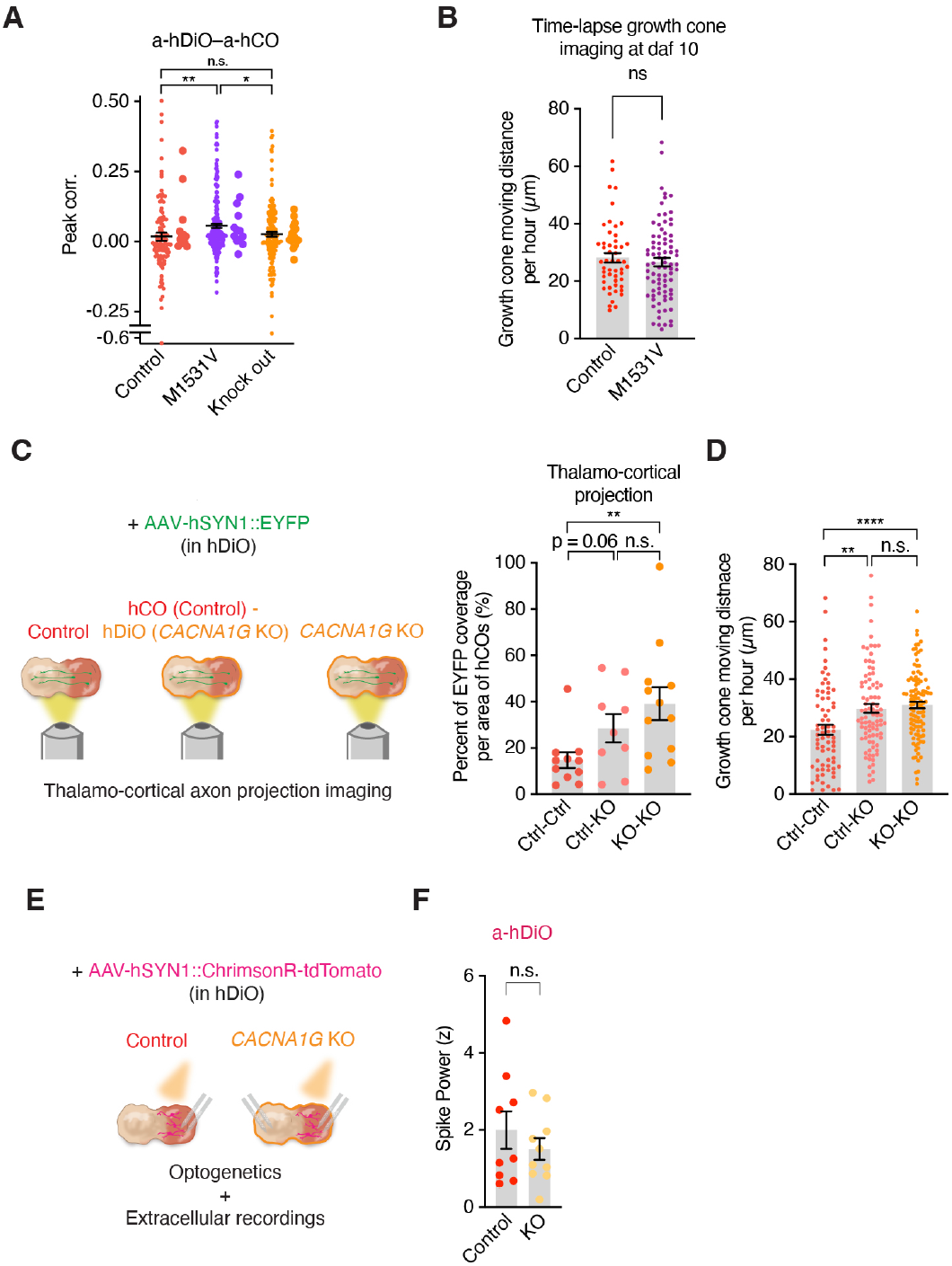
Gain- and loss-of-function *CACNA1G* variants in thalamo-cortical assembloids. (**A**) Scaled correlation analysis of unit pairs from a-hDiO to a-hCO from control, *CACNA1G* M1531V, and *CACNA1G* KO (n = 109 units from 12 assembloids, 9 differentiations for control, n = 201 units from 13 assembloids, 5 differentiations for *CACNA1G* M1531V and n = 159 units from 16 assembloids, 7 differentiations for *CACNA1G* KO; Kruskal-Wallis test, \**P* = 0.0406, \*\**P* = 0.0015) at daf 35–136. (**B**) Quantification of growth cone dynamic at daf 10 (n = 51 growth cones from 6 assembloids for control, n = 83 growth cones from 7 assembloids for *CACNA1G* M1531V; Mann-Whitney test, n.s., not significant). (**C**) Schematic illustrating thalamocortical projections in hCO–hDiO assembloids (left) and quantification (right) at daf 20 (*n* = 11 assembloids for control, n = 9 assembloids for Control–KO hybrid and n = 12 assembloids for *CACNA1G* KO; Kruskal-Wallis test, n.s., not significant, \*\**P* = 0.0050). (**D**) Quantification of growth cone dynamic at daf 10 (n = 73 growth cones from 8 assembloids for control, n = 90 growth cones from 6 assembloids for control–KO hybrid and n = 106 growth cones from 7 assembloids for *CACNA1G* KO; Kruskal-Wallis test, \*\**P* = 0.0013, \*\*\*\**P* < 0.0001). (**E**) Schematic illustration of the experimental design for hDiO optogenetic and extracellular recording (left). (**F**) Quantification of z-scored power of spike band frequency in a-hDiO during light stimulation (n = 9 assembloids for control and n = 10 assembloids for *CACNA1G* KO; Mann-Whitney test, n.s., not significant). Data is shown as mean ± s.e.m. Small dots indicate units or growth cones in A, B and large dots indicate assembloids in C, F.

## Notes

### Summary of Updates

new experiments, increased sample size and revised interpretation

