## Supplementary Tables 1 & 2 for "Human assembloids reveal the consequences of *CACNA1G* gene variants in the thalamocortical pathway" (from another version of this preprint: version 1, 2023-03-16)

**SUPPLEMENTARY TABLE 1.** hiPS cell lines used in this study.

| **Experiment / Line** | 1205-4 | 2242-1 | 0410-1 | 8119-3 | 1208-2 | KO#1 | KO#2 | M1531V |
|---|---|---|---|---|---|---|---|---|
| qPCR characterization | O | O | O | O | O | | | |
| Single cell RNAseq | O | | | O | O | | | |
| IHC characterization | O | | O | O | O | O | O | O |
| Axon projection | | O | O | | O | | O | O |
| Extracellular recording | O | | O | | O | O | O | O |
| Optogenetics + Calcium imaging | O | | | | O | | | |

**SUPPLEMENTARY TABLE 2.** Primers used for qPCR experiments.

| **Gene** | **Forward Primer** | **Reverse Primer** |
|---|---|---|
| *GAPDH* | CATGAGAAGTATGACAACAGCCT | AGTCCTTCCACGATACCAAAGT |
| *FOXG1* | AACCTGTGTTGCGCAAATGC | AAACACGGGCATATGACCAC |
| *TCF7L2* | AGAAACGAATCAAAACAGCTCCT | CGGGATTTGTCTCGGAAACTT |
| *OTX2* | CAAAGTGAGACCTGCCAAAAAGA | TGGACAAGGGATCTGACAGTG |
| *CACNA1G* | TGTCTCCGCACGGTCTGTAA | AAGCCGGTTCCAAGTGTCTC |
